# Chromosome-level genome assembly of *Euphorbia peplus*, a model system for plant latex, reveals that relative lack of Ty3 transposons contributed to its small genome size

**DOI:** 10.1101/2022.10.13.512124

**Authors:** Arielle R. Johnson, Yuanzheng Yue, Sarah B. Carey, Se Jin Park, Lars H. Kruse, Ashley Bao, Alex Harkess, Asher Pasha, Nicholas J. Provart, Gaurav D. Moghe, Margaret H. Frank

## Abstract

*Euphorbia peplus* (petty spurge) is a small, fast-growing plant that is native to Eurasia and has become a naturalized weed in North America and Australia. *E. peplus* is not only medicinally valuable, serving as a source for the skin cancer drug ingenol mebutate, but also has great potential as a model for latex production owing to its small size, ease of manipulation in the laboratory, and rapid reproductive cycle. To help establish *E. peplus* as a new model, we generated a 267.2 Mb HiC-anchored PacBio HiFi nuclear genome assembly with an embryophyta BUSCO score of 98.5%, a genome annotation based on RNA-seq data from six tissues, and publicly accessible tools including a genome browser and an interactive organ-specific expression atlas. Chromosome number is highly variable across *Euphorbia* species. Using a comparative analysis of our newly sequenced *E. peplus* genome with other Euphorbiaceae genomes, we show that variation in *Euphorbia* chromosome number is likely due to fragmentation and rearrangement rather than aneuploidy. Moreover, we found that the *E. peplus* genome is relatively compact compared to related members of the genus in part due to restricted expansion of the Ty3 transposon family. Finally, we identify a large gene cluster that contains many previously identified enzymes in the putative ingenol mebutate biosynthesis pathway, along with additional gene candidates for this biosynthetic pathway. The genomic resources we have created for *E. peplus* will help advance research on latex production and ingenol mebutate biosynthesis in the commercially important Euphorbiaceae family.

**Significance statement:** *Euphorbia* is one of the five largest genera in the plant kingdom. Despite an impressive phenotypic and metabolic diversity in this genus, only one *Euphorbia* genome has been sequenced so far, restricting insights into *Euphorbia* biology. *Euphorbia peplus* has excellent potential as a model species due to its latex production, fast growth rate and production of the anticancer drug ingenol mebutate. Here, we present a chromosome-level *E. peplus* genome assembly and publicly accessible resources to support molecular research for this unique species and the broader genus. We also provide an explanation of one reason the genome is so small, and identify more candidate genes for the anticancer drug and related compounds.

## Introduction

The Euphorbiaceae is a large plant family with over 6,000 species. Almost all Euphorbiaceae species produce a milky terpenoid-rich substance called latex, which is contained in specialized cells and exudes from damaged tissue. Euphorbiaceae latex is used for producing natural rubber (e.g. *Hevea brasiliensis*, Pará rubber tree)(Yamashita & Takahashi 2020), is a carbon source for biofuels (e.g. *Euphorbia lathyris*, caper spurge)(Pan et al. 2022), and contains unique biosynthetic pathways that support the production of medically-relevant compounds(Yang Xu et al. 2021). Latex from *Euphorbia peplus* (commonly known as ‘petty spurge’ and ‘cancer weed’) contains a diterpenoid compound called ingenol mebutate that is used in pharmaceutical treatments for skin cancer(Lebwohl et al. 2012), making this species particularly valuable. While parts of the biosynthetic pathway for ingenol mebutate have recently been identified(Czechowski et al. 2022), the full pathway has yet to be elucidated, and pharmaceutical production is currently limited to natural extraction from *E. peplus*.

Several economically valuable Euphorbiaceae crop plants have extensive genome resources, including *Hevea brasiliensis* (Pará rubber tree), *Manihot esculenta* (cassava), *Jatropha curcas* (physic nut), and *Ricinus communis* (castor oil plant), but these are physically large crop species that are not ideal for laboratory work. The cells that produce and contain latex, laticifers, are not developmentally well characterized; understanding this network of living tubes will require a model species that is easy to manipulate (Johnson et al. 2021). A smaller model species that can be grown in the lab, *Euphorbia lathyris* (caper spurge), was developed as a model system for latex in the Euphorbiaceae, and the generation of *Euphorbia lathyris* mutants that produce more or less latex than wild-type plants has produced insights into latex development(Castelblanque et al. 2016, 2018, 2021). The *Euphorbia lathyris* genome was also recently published(Mingcheng Wang et al. 2021). However, *Euphorbia lathyris* is not suitable for some experiments because of its biennial life cycle and large stature (>1 meter at maturity).

We have developed genomic resources for *E. peplus* as a complementary model system to *Euphorbia lathyris* to study latex development in the Euphorbiaceae. *E. peplus* and *E. lathyris* are in the same subgenus, *Esula*; *E. peplus* is in section *Tithymalus* while *E. lathyris* is in section *Lathyris*, the earliest diverging section within *Esula*, so the two species’ lineages diverged approximately 40 million years ago(Riina et al. 2013; Anest et al. 2021). *E. peplus* has a relatively small genome size of <300 Mb, is an annual plant with a short life cycle of ∼6 weeks post-germination to flowering, and is only ∼30 cm tall at maturity, allowing it to be grown in relatively large quantities in growth chambers (**Figure 1**). Virus-induced gene silencing (VIGS) has been successfully performed in this species(Czechowski et al. 2022), making it a good candidate to identify and functionally test developmental and biochemical genes of interest.

**Figure 1:**
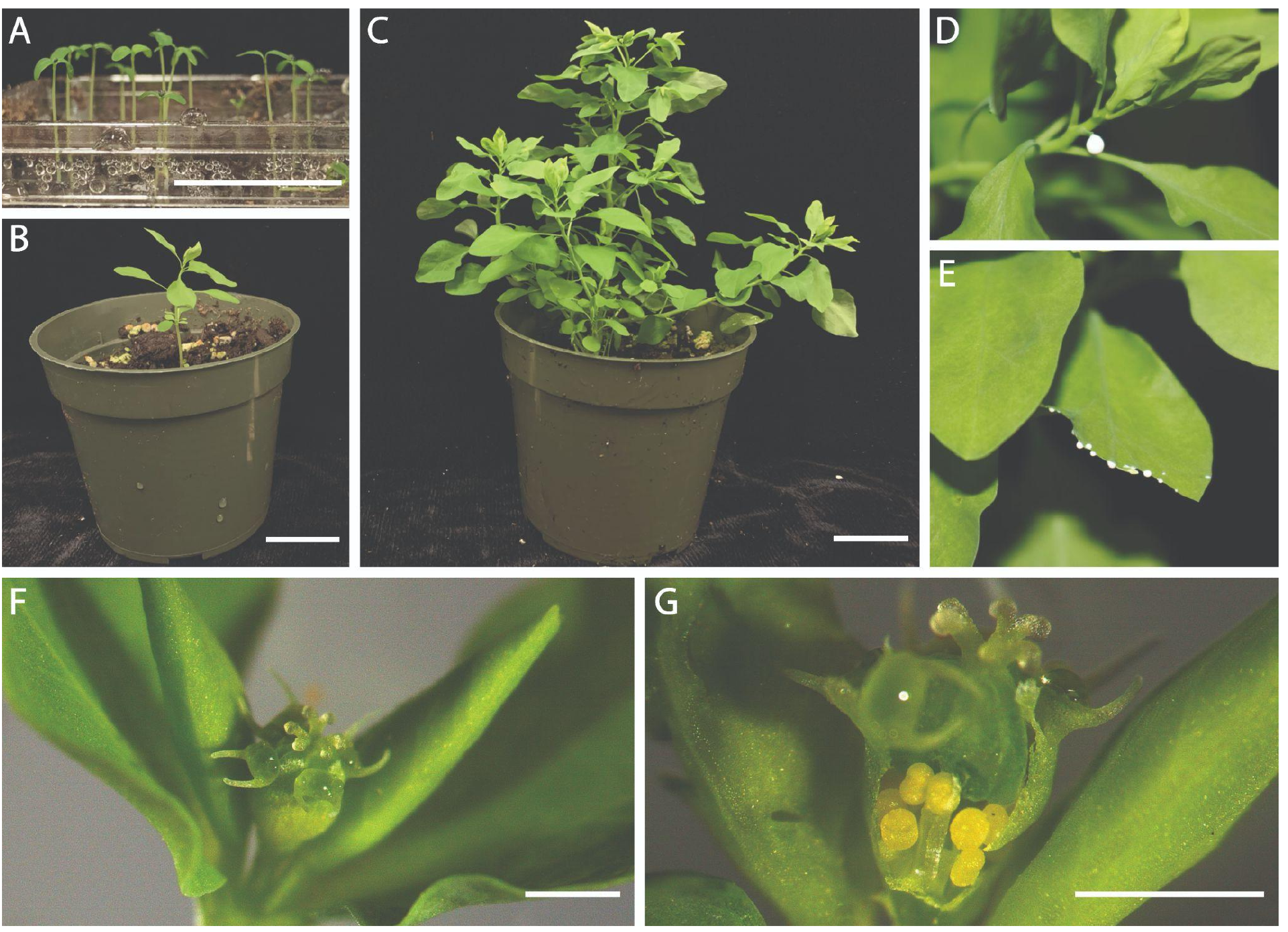
*Euphorbia peplus* is a small, rapidly-maturing plant that produces latex. Scale in A, B, and C is 30mm; scale in F and G is 1mm. A. 1-week-old seedlings; lid of tray removed for photo. B. 3-week-old plant with vegetative growth only. C. 5.5-week-old plant that is reproductively mature. D. Cut petiole exuding latex. E. Cut leaf exuding latex. F. Exterior view of inflorescence showing stigmas and nectaries; rolled bracts to the left and right conceal developing inflorescences. G. Inflorescence is cut to reveal staminate flowers.

This paper presents the nuclear genome of *E. peplus*, puts the genome into an evolutionary context using other published Euphorbiaceae genomes, examines why the *E. peplus* genome is uniquely small, and provides new hypotheses regarding the evolution of a valuable diterpene biosynthetic pathway in this species. In addition, introduction of this new genome expands the resources available for genomic studies of the phenotypically diverse *Euphorbia* genus.

## Results

### *A chromosome-scale assembly of the* Euphorbia peplus *genome*

To build a nuclear genome assembly for *E. peplus*, we first generated 22.7 Gb of PacBio HiFi Circular Consensus Sequence (CCS) reads and 48.4 Gb of paired-end 150 nt Phase Genomics Hi-C reads. K-mer based analysis of the raw HiFi reads suggests that the genome size of the accession is 252.2 Mb (**Supplementary Figure 1**) and that the genome is highly homozygous (99.9%), consistent with the fact that *E. peplus* is a self-compatible plant which typically self-fertilizes(Asenbaum et al. 2021). An initial PacBio HiFi assembly resulted in a nuclear assembly size of 327.6 Mb assembled into 1210 contigs with a contig N50 of 23.9 Mb and a L50 count of n=6 contigs. After Hi-C scaffolding and manual editing, our final assembly comprises 330.5 Mb assembled into 1242 scaffolds with a scaffold N50 of 31.0 Mb and L50 count of n=5 scaffolds (**Supplementary Table 1**). After selecting the chromosomes only, our final chromosomal assembly consists of 8 chromosomes of >20Mb each, totaling 267.2 Mb, in agreement with previous chromosome squashes and previous flow cytometry analyses(Fasihi et al.; Loureiro et al. 2007) (**Figure 2A**). Benchmarking of universal, single-copy orthologs (BUSCO) analysis of those chromosomes indicated a mostly complete assembly with 98.5% of Embryophyta BUSCO orthologs identified, most of which (95.5%) were single-copy (**Supplementary Table 2**). The chromosomal genome assembly size, 267.2 Mb, is only 6% larger than our 21-mer genome size estimate, 252.2 Mb, and the chromosomes constitute a 98.9% complete genome according to a Merqury kmer completeness analysis (**Supplementary Table 3**). The 63.3 Mb of non-chromosome scaffolds do not contain any BUSCO orthologs, most of the annotated genes appear to be chloroplastic based on their human-readable descriptions, the Extensive de novo TE Annotator (EDTA) only annotated 0.97% of the sequences as repetitive elements (**Supplementary Table 4**), and alignment of the raw Hi-C data against the non-chromosomal scaffolds only results in 3.52% uniquely mapped reads compared with 42.20% uniquely mapped reads when mapped against the chromosomes. This combination of evidence leads us to conclude that the non-chromosomal scaffolds are mostly not nuclear DNA, and the chromosomal assembly is likely close to the actual genome size.

**Figure 2:**
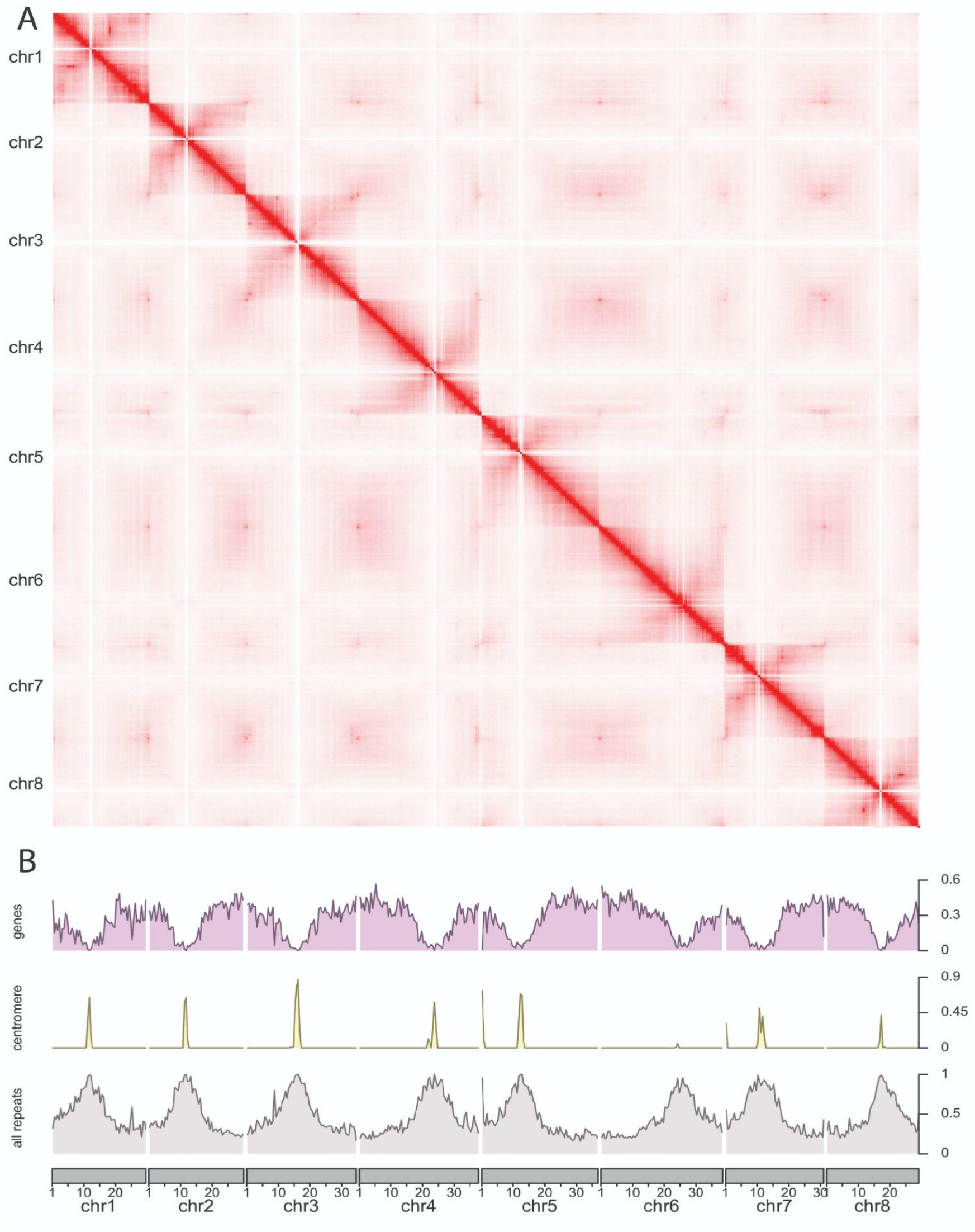
The *E. peplus* reference genome is a high-quality, chromosome-scale assembly. A. Hi-C contact map of the 8 chromosomes assembled for the *E. peplus* genome generated with Juicebox. B. Coverage of annotated genes, putative centromeric repeats, and all masked repeats.

### *The* Euphorbia peplus *genome annotation includes human-readable descriptions and GO terms*

We masked repeats in the genome using RepeatModeler and RepeatMasker, which led to masking of 57.66% of the nucleotides in all scaffolds and 48.55% of the nucleotides in the assembled chromosomes. The most common retroelements by far were Ty1/Copia, comprising 14.73% of the chromosomes, and Ty3, comprising 4.76% of the chromosomes. (Ty3 is the family previously referred to as “Gypsy” in some publications; the transposon family name has been reconsidered because it is insensitive to people of Romani heritage(Wei et al. 2022).) The most common DNA transposon, Harbinger, comprised 0.31% of the chromosomes. We then used the BRAKER pipeline to predict protein coding genes using both homology with proteins from the OrthoDB v10 plant database and short-read RNAseq evidence from six tissue types (**Supplementary Table 5, Supplementary Figure 2**). This analysis generated 27,228 total gene annotations: 25,471 primary gene transcripts and 1,757 alternate transcripts. For these gene models, 99.0% of Embryophyta BUSCO orthologs were identified and 90.1% were single-copy (**Supplementary Table 6**). Next, we used Automatic assignment of Human Readable Descriptions (AHRD) to produce 20,929 human-readable gene names for these annotations, 17,639 of which were highest-quality. Using BLAST2GO, we assigned functional labels to 84% of the annotations. A partial centromeric repeat was determined by taking the top result from Tandem Repeats Finder. As expected, we found that gene density declines near the centromere locations of the chromosomes (**Figure 2B**). We also created a genome browser using the JBrowse platform and an interactive expression atlas using the eFP browser to make the genome annotation readily accessible online(Buels et al. 2016; Sullivan et al. 2019).

### *Differences in genome architecture between* E. peplus *and* E. lathyris *are not due to aneuploidy*

The genus *Euphorbia* has a highly variable chromosome count, with base numbers ranging from n=6 to n=10,(Wurdack et al. 2005) despite the fact that there is no evidence for a whole-genome duplication event in the *Euphorbia* lineage(Mingcheng Wang et al. 2021). One hypothesis is that this variation was driven by aneuploidy (i.e. offspring inheriting an extra chromosome or a missing chromosome)(Hans 1973). To investigate the conservation of chromosomal architecture, we visualized the macrosynteny between the *E. peplus* genome and the other publicly-available chromosome-level Euphorbiaceae genome assemblies, namely *Euphorbia lathyris, Manihot esculenta, Hevea brasiliensis*, and *Ricinus communis*.

Based on our results, aneuploidy is not responsible for the difference in chromosome number between the 8 chromosomes in *E. peplus* and the 10 chromosomes in *E. lathyris*. If chromosomes had been duplicated, we would observe macrosynteny between an entire *E. peplus* chromosome and multiple entire *E. lathyris* chromosomes. Instead, multiple chromosomal fragmentation and rearrangement events seem to have contributed to the difference in chromosome number (**Figure 3**). For example, large parts of *E. peplus* chromosome 6 are homologous to large parts of *E. lathyris* chromosome 2 and *E. lathyris* chromosome 5, but *E. lathyris* chromosome 2 also has large parts that are homologous to parts of *E. peplus* chromosome 3 and *E. lathyris* chromosome 5 also has large parts that are homologous to parts of *E. peplus* chromosome 5. This suggests that the chromosomes were most likely fragmented rather than duplicated.

**Figure 3:**
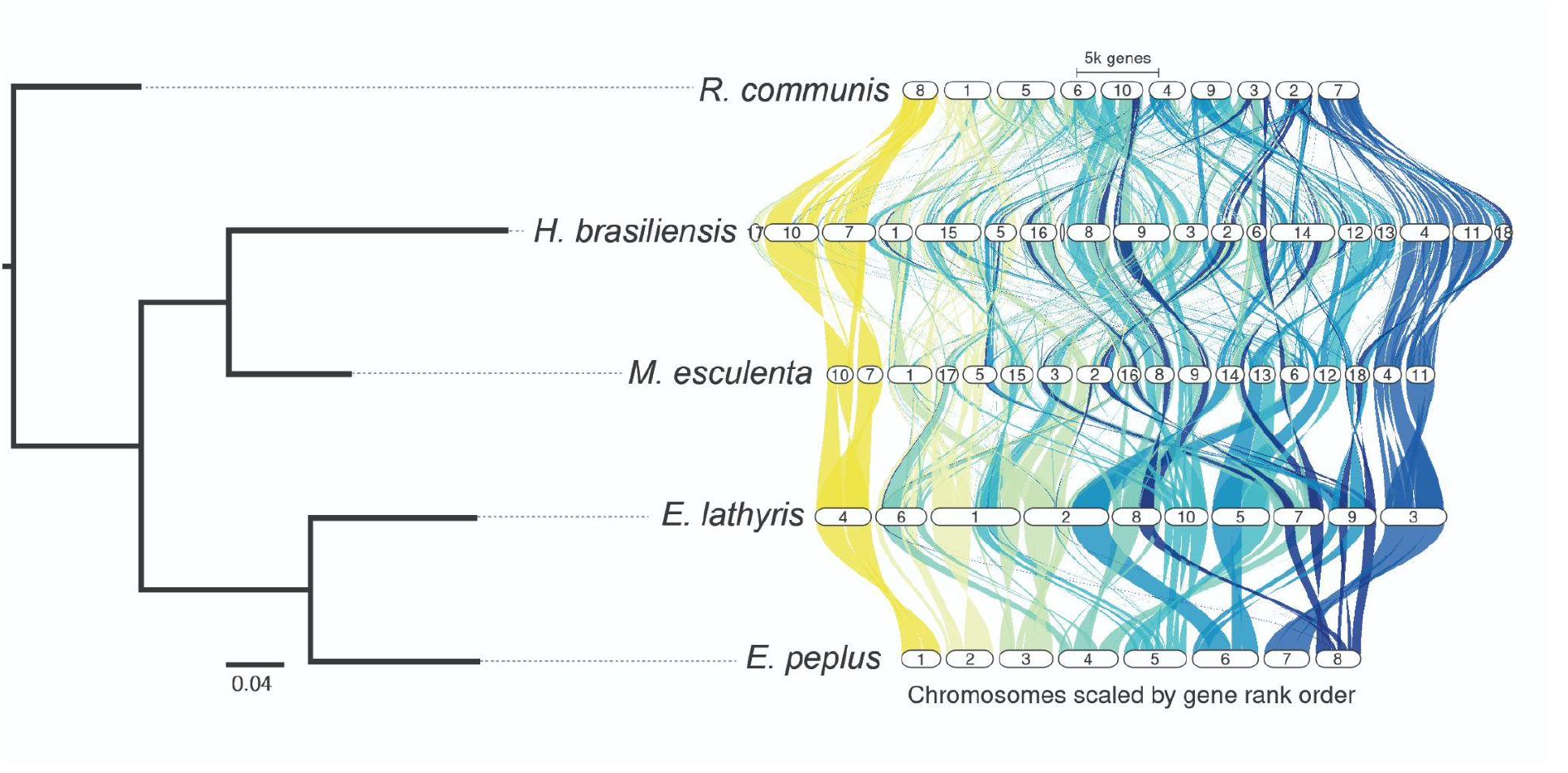
Evidence of chromosomal fragmentation and rearrangement within *Euphorbia*. Left: Species phylogeny produced by OrthoFinder. Right: GENESPACE plot of *E. peplus* chromosomal synteny with other Euphorbiaceae species that have publicly available chromosome-level genome assemblies. Colors are used to represent independent chromosomes in the *E. peplus* genome.

### *Difference in genome size between* E. peplus *and* E. lathyris *is largely explained by relative lack of TEs, especially Ty3, in* E. peplus

Differences in plant genome size are thought to arise largely through differential accumulation of transposable elements (TEs) and through whole genome duplication events(Michael 2014; Dandan Wang et al. 2021). Therefore, in order to investigate why the *E. peplus* genome is so small compared to that of *E. lathyris* despite no evidence of recent whole-genome duplication in *E. lathyris*, we compared the TE composition between the *E. peplus* genome and the other available chromosome-level Euphorbiaceae genomes. For the five Euphorbiaceae species, the species with the larger chromosomal genomes also generally had a higher proportion of repetitive elements in their genome, supporting the idea that TE content is largely responsible for fluctuations in genome size for this family (**Figure 4A**). Compared with the other species, *E. peplus* has a much lower proportion of Ty3 TEs (**Figure 4B**): For example, *E. peplus* has 12.7Mb of Ty3 sequence whereas *E. lathyris* has 205.5Mb of Ty3 sequence, over 16x as much Ty3 sequence as *E. peplus*. The most parsimonious explanation for this observation is that Ty3 elements have remained suppressed over time in *E. peplus*, although sequencing more taxa and inferring the ancestral state of the *Esula* subgenus may be required to detect the actual mechanisms leading to genome size differences. Nonetheless, this lack of Ty3 accounts for the most substantial difference in overall TE abundance between *E. peplus* and the other Euphorbiaceae genomes. A difference for Copia also exists but is less stark. *E. peplus* has 39.3Mb of Copia sequence, while *E. lathyris* has 253.1Mb of Copia sequence, around 6.5 times as much Copia sequence as *E. peplus. E. peplus*’ total repetitive sequences are 129.7Mb while the total in *E. lathyris* is 766.9Mb, while the non-repetitive content is 137.5Mb in *E. peplus* and 219.9Mb in *E. lathyris*. Put differently, if *E. peplus* had the same absolute quantity of repetitive sequences as *E. lathyris* added to its existing non-repetitive sequences, its chromosomes would be 904.4Mb (close to *E. lathyris*’ 986.8Mb) rather than 267.2Mb, again suggesting that TEs make up most of the difference in genome size between these two species.

**Figure 4:**
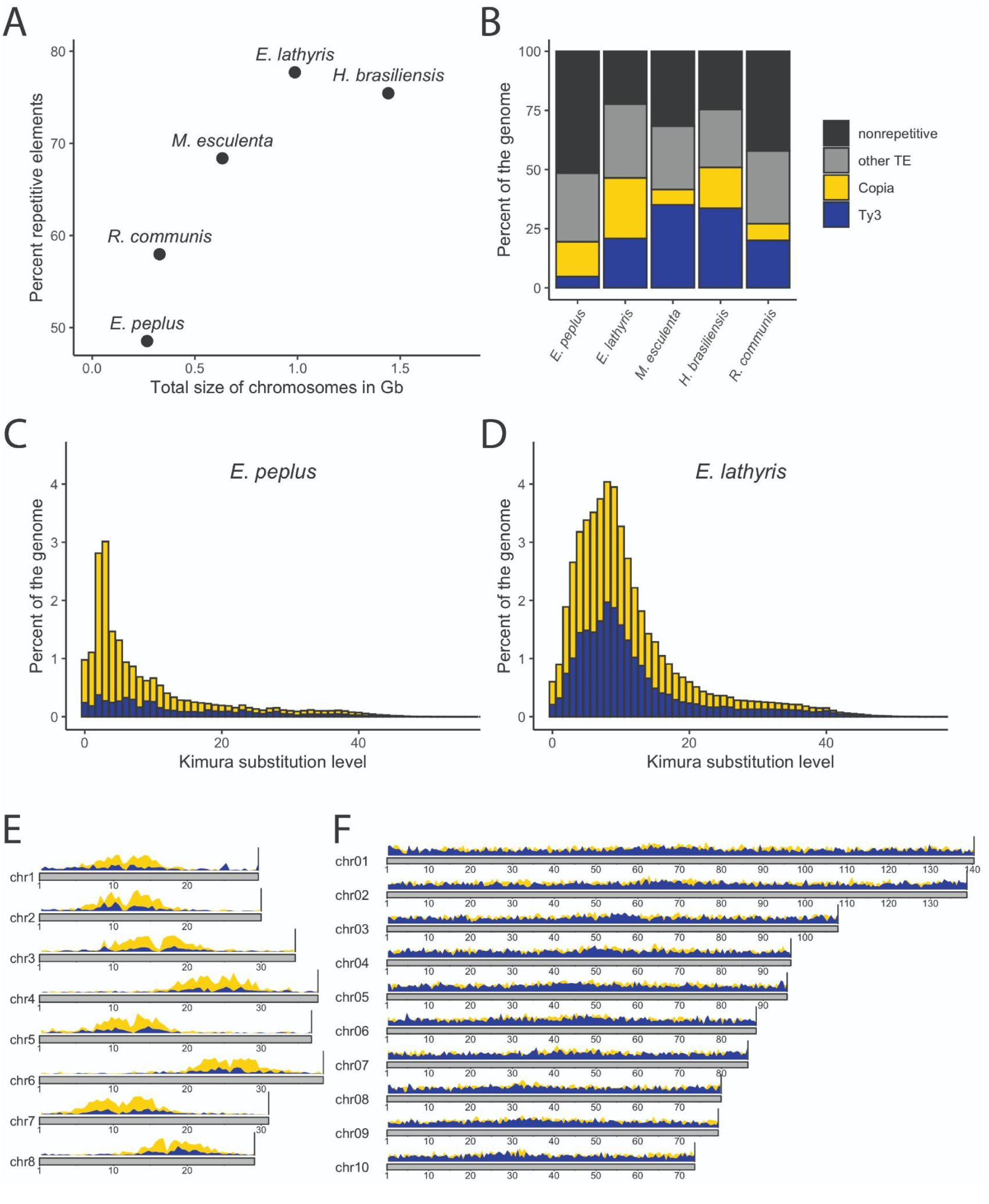
*E. peplus* has a low Ty3 copy number compared to related species. A. Scatterplot of total size of all chromosomes in each genome versus the percent of repetitive elements in the chromosomes. B. Percentages of all chromosomes masked for Ty3, Copia, and all other TEs. C. Stacked Kimura distance plot for *E. peplus* showing Ty3 (dark blue) and Copia (yellow). D. Stacked Kimura distance plot for *E. lathyris* showing Ty3 (dark blue) and Copia (yellow). E. KaryoPloteR density plot of Ty3 (dark blue) and Copia (yellow) across *E. peplus* chromosomes; y-axis bar to right of each plot is 0.7. F. Coverage plot of Ty3 (dark blue) and Copia (yellow) across *E. lathyris* chromosomes; y-axis bar to right of each plot is 0.7.

We also examined the age of the Copia and Ty3 repeats in the *E. peplus* and *E. lathyris* genomes by calculating the Kimura distance: the number of substitutions between each instance of a certain repeat in the genome and that repeat family’s consensus sequence (an approximation of the ancestral progenitor’s sequence, created by taking the most common nucleotide at each site across a multiple alignment of the copies of the repeat). Kimura distance serves as a proxy for the history of the expansion of TE families, as younger elements are expected to be similar to the consensus sequences whereas older elements are thought to have accumulated more mutations over time (Kimura 1980). It appears that there have been no major changes in Ty3 abundance over evolutionary time in *E. peplus*, whereas a substantially increased abundance of Ty3 elements is seen in *E. lathyris*, with its maximum at a Kimura substitution level of 8 (**Figure 4C and 4D**). Both *E. peplus* and *E. lathyris* have a peak of Copia expansion, but the *E. lathyris* peak is older, with a maximum at a Kimura substitution level of 8, whereas the *E. peplus* peak is much younger and narrower and has a maximum at a Kimura substitution level of 3. Note that *E. peplus* has a shorter generation time than *E. lathyris*, as *E. peplus* is an annual plant while *E. lathyris* is biennial; however, shorter generation time would theoretically lead to the accumulation of more mutations so our observation that the *E. peplus* peak is younger remains valid. We also examined the distribution of Ty3 and Copia TEs abundance across the length of the chromosomes in *E. peplus* and *E. lathyris*. Interestingly, both Ty3 and Copia decrease in abundance with distance from the centromere more drastically in *E. peplus* than they do in *E. lathyris* (**Figure 4E and 4F**); however, the significance of this differential distribution is not clear.

RNA polymerase V subunit NRPE1 has been shown to regulate TE abundance in Arabidopsis. NRPE1 is the largest subunit of plant RNA polymerase V, which helps produce the RNA scaffolds necessary for the RNA-directed DNA methylation that causes transgenerational suppression of TE activity(Matzke & Mosher 2014). NRPE1 has been identified through genome wide association studies (GWAS) as a major determinant of genome-wide CHH methylation patterns and specifically, of CHH methylation of TEs in Arabidopsis(Sasaki et al. 2019). It is plausible that a duplicate of NRPE1 could be functional in *E. peplus* given that a similar example of NRPE1 duplication-neofunctionalization exists in the grasses (Poaceae)(Trujillo et al. 2018). To examine a potential role for NRPE1 in the regulation of Ty3 copy number in *E. peplus*, we produced orthologous groups containing proteins from *E. peplus* and the other four Euphorbiaceae genomes used in this paper (**Supplementary Figure 3**), and examined the orthologous groups containing genes that are known to regulate TE abundance (**Supplementary Table 7, Supplementary File 4**). In our results, the NRPE1 orthogroup showed a *E. peplus*-specific duplication and was not duplicated in any other species, although one of its two *E. peplus* copies was broken into multiple gene models, indicating that it may be a pseudogene; low levels of expression were present (**Supplementary Figure 4-6**). Whether the additional NRPE1 copy in *E. peplus* is, or was, functional and could explain the suppression of Ty3 requires further investigation.

### Diterpenoid biosynthetic gene candidates previously thought to be localized to two clusters actually constitute a single cluster

Elucidating the biosynthetic pathway of ingenol mebutate and other diterpenoids of medical relevance is a research priority in *Euphorbia(Bergman et al. 2019; Ricigliano et al. 2020; Forestier et al. 2021)*. Ingenol mebutate has a 5/7/7/3 carbon ring system. In the first step of the proposed pathway for ingenol mebutate biosynthesis, casbene synthase cyclizes geranylgeranyl diphosphate and removes its phosphate groups, producing casbene, which contains the 3-carbon ring(Luo et al. 2016). Then multiple Cytochrome P450 (CYP450) enzymes add three hydroxyl groups to the molecule, and the dehydrogenation of those hydroxyl groups by an alcohol dehydrogenase sets off a spontaneous intramolecular aldol reaction that forms the 5-carbon ring and produces the intermediate Jolkinol C (Wong et al. 2018; Forestier et al. 2021). The subsequent steps that convert Jolkinol C to ingenanes including ingenol mebutate are not currently clear (**Supplementary Figure 7**). A recent publication described two putative diterpenoid biosynthetic gene clusters in *E. peplus* based on a bacterial artificial chromosome library(Czechowski et al. 2022). One cluster contained casbene synthase, casbene 5-oxidase (CYP726A19) and an alcohol dehydrogenase, and the other contained casbene 5-oxidase (CYP726A4) and casbene-9-oxidase (CYP71D365). Based on our genome assembly and annotation, these are actually one contiguous biosynthetic gene cluster spanning ∼0.6 Mb, containing most of the currently functionally characterized members of the pathway (**Figure 5, Supplementary Table 8**). The only functionally characterized putative ingenol mebutate biosynthetic gene that is not present in this cluster is the specific alcohol dehydrogenase functionally characterized in Luo et al 2016; its top BLAST hit in our genome is on another chromosome. However, given that in yeast, using an *E. lathyris* alcohol dehydrogenase versus a *Jatropha curcas* alcohol dehydrogenase does not produce statistically different levels of Jolkinol C(Wong et al. 2018), it is possible that a different enzyme could be used interchangeably in this pathway.

**Figure 5:**
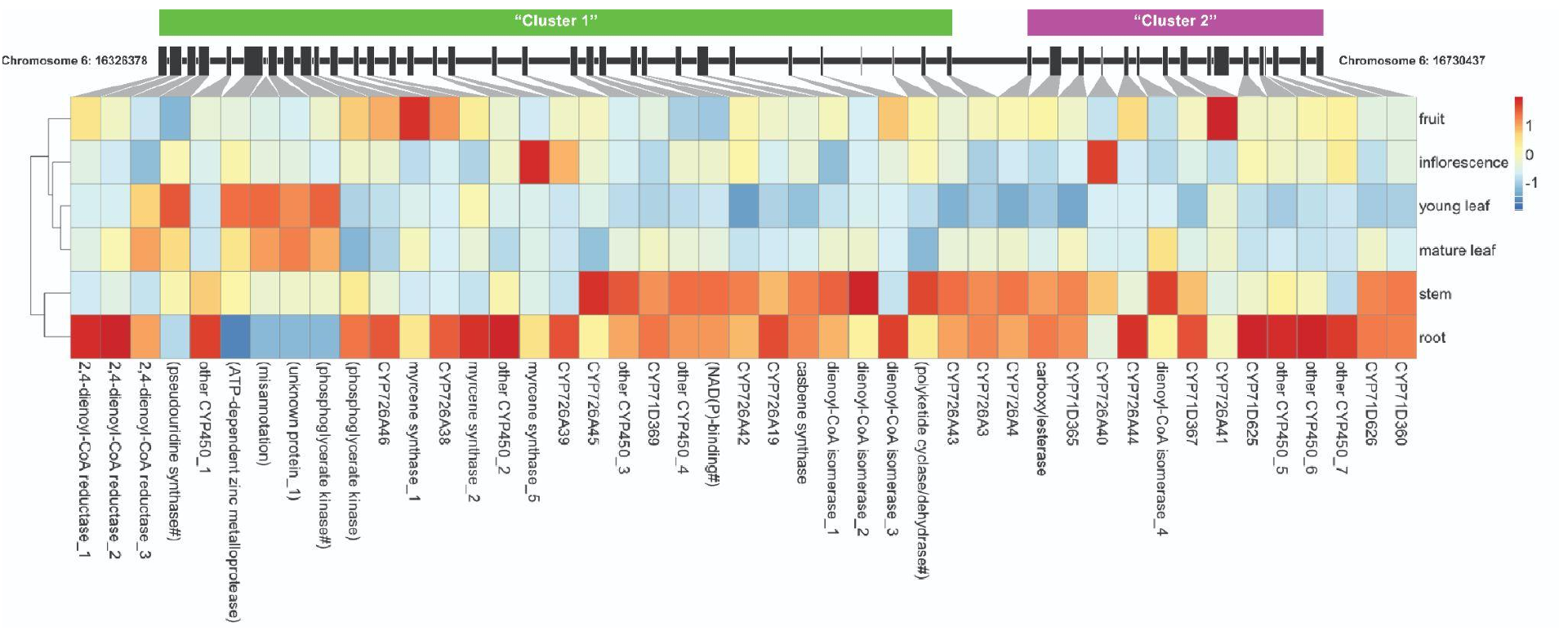
A diterpenoid biosynthetic region previously reported to be two clusters is actually a single cluster. The genes’ location is shown along the chromosome, and the cell color corresponds to the expression data averaged by sample and normalized across tissues for each gene. Genes with raw expression counts of less than 20 are not shown in the heatmap. Cluster names and Cytochrome P450 (CYP450) names are following Czechowski et al 2022. Genes unlikely to be involved in diterpenoid biosynthesis are listed in parentheses. Genes with long gene names are abbreviated with a “#”, and full gene names are in **Supplementary Table 8**.

## Discussion

This paper introduces a new high-quality nuclear genome assembly and annotation for *E. peplus*, and examines the evolution of its karyotype, TE landscape, and diterpenoid biosynthesis. Based on the two currently available *Euphorbia* genomes, chromosome fragmentation and fusion appear to drive chromosome count variation in the genus. As more Euphorbiaceae and *Euphorbia* genomes are released, especially those of species that diverged more recently in evolutionary time, we will get closer to understanding the evolutionary history of this genus and why its karyotype varies so dramatically. It would be interesting to investigate whether chromosome fragmentation events are adaptively neutral or whether they have helped enable innovations such as the repeated evolution of carbon concentrating mechanisms that have allowed *Euphorbia* species to occupy an extremely wide range of habitats(Horn et al. 2014).

Our analysis shows that ∼89% of the size difference between the *E. peplus* and *E. lathyris* genomes could be due to differences in TE abundance, in agreement with other recent studies that have emphasized the importance of TEs in genome size evolution(Dandan Wang et al. 2021; Akakpo et al. 2020). Differential accumulation of Ty3 specifically has been shown to affect genome size in other species — for example, in the Brassicaceae, Ty3 is linked to the increased genome size in *Arabis alpina* compared with *Arabidopsis thaliana(Willing et al. 2015)*. Moreover, studies in *Arabidopsis thaliana* show that Ty3 transposons were more frequently deleted than other classes of transposons across a panel of 216 *Arabidopsis thaliana* accessions(Stuart et al. 2016), suggesting genetic lability of the Ty3 family in particular. Further research examining the role of CHH methylation in suppressing Ty3 propagation and the propagation of other TEs may help explain why *E. peplus* has retained a relatively compact genome. Toward that end, the extra NRPE1 paralog in *E. peplus* could be further investigated; the fact that Copia TEs peaked only recently in *E. peplus* could be consistent with a recent pseudogenization event of a previously functional NRPE1 homolog. It would also be interesting to investigate whether *E. peplus*’ mating system plays a role, as *E. peplus*’ inflorescences self-fertilize whereas *E. lathyris* requires pollinators. Mathematical modeling predicts a lower abundance of TEs in populations with more selfing because selfing makes it more difficult for new TE copies to invade the genome and be transmitted at non-Mendelian frequencies(Boutin et al. 2012); however, in empirical studies TE dynamics have been shown to vary by species in ways that do not simply reflect their mating strategy(Agren et al. 2014; Legrand et al. 2019).

In this paper we show that many of the putative biosynthetic pathway genes for important diterpenoids are highly expressed in *E. peplus* stems and roots, which is where *Euphorbia* diterpenoids are generally concentrated(Ernst et al. 2019). However, ingenol mebutate itself is most abundant in the stem latex, not in the roots(Czechowski et al. 2022). Perhaps the biosynthesis of ingenol mebutate takes place across multiple cell types, as with morphine biosynthesis in opium poppies where only the final steps occur in laticifers (the cells that contain latex)(Onoyovwe et al. 2013). It is also interesting that so many of the putative diterpenoid biosynthetic genes form a cohesive cluster. Plant biosynthetic gene clusters for specialized metabolites have been found across multiple species(Nützmann et al. 2018; Polturak & Osbourn 2021). A diterpenoid cluster in the Euphorbiaceae had previously been hypothesized and detected by long-distance PCR(King et al. 2014); this paper corroborates those previous findings and provides the greatest evidence to date of a diterpenoid biosynthetic cluster in this family. In addition to functionally characterizing more candidate biosynthetic genes, future studies could examine whether the tight clustering of putative diterpenoid biosynthetic genes easily enables the plant to regulate transcription of that pathway, or whether the clustering enables the biosynthetic enzymes to form a metabolon, a noncovalent interaction of sequential enzymes in a pathway in physical space that functions like an “assembly line”(Nützmann et al. 2016).

In conclusion, our *E. peplus* genome assembly provides insights into the large variation in chromosome number, variation in genome size, and unique chemistry of the Euphorbiaceae. This genome resource will be useful for identifying candidate genes for further elucidation of diterpenoid biosynthesis in this species, including the synthesis of ingenol mebutate and other compounds of clinical relevance. Moreover, this assembly makes *E. peplus* an ideal complementary model system to *Euphorbia lathyris* for studying latex development in the Euphorbiaceae. We are releasing our assembly with web-accessible tools including a genome browser and interactive expression atlas. This assembly is especially important for the study of the *Euphorbia* genus, given that it is one of the largest genera of flowering plants with >2000 species are documented(Esser et al. 2009), and only one other chromosome-level genome has been published. It is our goal to advance our understanding of the unique evolutionary and metabolic biology of the Euphorbiaceae by releasing this high-quality genomic resource.

## Materials and methods

### Plant materials and growth conditions

*E. peplus* plants were obtained from the Cornell Botanical Gardens area: seeds set by the wild *E. peplus* plants were collected. The taxonomic identity of *E. peplus* was verified by sequencing a diagnostic region of the *matK* gene using the following primers (Forward: 5’-CCC CAT CCA TCT CGA AAA ATT GG-3’; Reverse: 5’-ATA CGC GCA AAT TGG TCG AT-3’), and through morphological characterization. The *matK* sequences obtained via Sanger sequencing **(Supplementary File 1)** were validated as belonging to *E. peplus* using the NCBI blastn tool, using the nr database as query. For initial DNA and RNA experiments, *E. peplus* seeds were cleaned using 10% trisodium phosphate solution and germinated on filter paper in petri dishes, moistened with 1 ml of 200uM Gibberellic acid 3 to promote germination. Organs for genome sequencing and RNA-seq were sampled from mature flowering stage plants. For subsequent experiments including imaging, seeds were germinated without pretreatment in closed Phytatrays (P5929,Sigma-Aldrich, Saint Louis, MO, U.S.A.) in Cornell soil mix(-Boodley & Sheldrake) or LM 1-1-1 soil mix. Plants were then transplanted to 4-inch pots containing Cornell soil mix or LM 1-1-1 soil mix and grown under long day conditions in 25 C day/16 C night conditions until flowering. Seeds were collected by placing mature plants in 12×16 organza party favor bags (QIANHAILIZZ, Amazon.com) and allowing them to dry for 3-4 days; plants were then composted and the contents of the bags collected. Seeds were then separated from chaff by rolling bag contents on a sheet of paper (round seeds are more mobile than chaff).

### Organ harvesting, RNA extraction, Illumina RNA-seq library prep, and sequencing

To generate an expression atlas and annotate the *E. peplus* genome, three biological replicates of the following organs were harvested from mature *E. peplus* plants: immature fruit, flowers, whole root, stem, young leaves, and mature leaves. Organs were flash frozen in liquid nitrogen immediately after harvesting, ground into a fine powder, and then processed for RNA extraction using a combined TRI Reagent (Sigma-Aldrich, Saint Louis, MO, U.S.A.) and Monarch Total RNA Miniprep Kit (New England Biolabs, Ipswich, MA, U.S.A.) protocol. One ml of TRI Reagent was added to approximately 50 mg of ground tissue, the samples were vortexed for 30 seconds, and then 200 µl of Chloroform was added. The samples were vortexed 3 × 30 seconds, left to sit at room temperature (RT) for 5 minutes, centrifuged at 12,000 G for 15 minutes at 4 ºC, and then the aqueous phase (the top layer) was transferred into a one-to-one mix with 25 Phenol:24 Chloroform:1 Isoamyl Alcohol (77617, Sigma-Aldrich, Saint Louis, MO, U.S.A.). The samples were vortexed for 30 seconds, and then centrifuged at 21,000 G for 10 minutes at 4 ºC. The top layer was transferred directly onto gDNA removal columns provided by the NEB Monarch Total RNA Miniprep Kit, and manufacturer guidelines were followed for Part 2 (RNA binding and elution) of the Monarch prep kit. RNA quality and quantity were accessed using a DeNovix DS-11 FX+ spectrophotometer (DeNovix Inc., Wilmington, DE, U.S.A.).

To construct RNA-seq libraries for Illumina sequencing, mRNA was isolated from 1,000 ng of total RNA using a NEBNext Poly(A) mRNA Isolation Module (E7490, New Englad Biolabs, Ipswich, MA, U.S.A.), followed directly by library construction using a NEBNext Ultra Directional RNA Library Prep Kit for Illumina (E7420). The libraries were barcoded with NEBNext Multiplex Oligos for Illumina Set 1 #E7335. The libraries were submitted to the Cornell Institute for Biotechnology Genomics Center, where they were quantified using qRT-PCR, quality checked on a Bioanalyzer (Agilent, Santa Clara, CA, U.S.A.), and pooled in equimolar ratios for 12-plex sequencing on a NextSeq 500 (Illumina, Hayward, CA, U.S.A.) 2×150 paired-end run.

### RNAseq read processing

In order to improve RNAseq data quality, raw RNAseq data was assessed for quality with FastQC(Andrews & Others 2010). RNAseq data was trimmed with Trimmomatic using the parameters SLIDINGWINDOW:5:20 MINLEN:90(Bolger et al. 2014). The RNAseq data was aligned to the genome assembly using STAR using its basic 2-pass mapping mode and default parameters(Dobin et al. 2013).

### DNA extraction and sequencing

In order to produce accurate long reads, plant tissue was ground in liquid nitrogen with a mortar and pestle and transferred to 2mL microcentrifuge tubes. A cetyl trimethylammonium bromide (CTAB) extraction method using chloroform:isoamyl alcohol 24:1, including treatment with Proteinase K and RNAse A, was used to extract the DNA(Fulton et al. 1995). DNA samples were sent to HudsonAlpha Institute for Biotechnology for PacBio HiFi circular consensus sequencing (CCS), where they were sheared with a Diagenode Megaruptor, size selected for 18kb fragments on the SageELF electrophoresis system and sequenced on a PacBio Sequel-II sequencer. A total of 1.3 million filtered CCS reads were generated, spanning 22.7 Gb or ∼90x genome coverage (based on the GenomeScope kmer genome size estimate of 252.2 Mb).

### HiC protocol and sequencing

In order to generate proximity ligation data, genomic DNA for HiC sequencing was crosslinked, fragmented, and purified from young leaf tissue from the same original plant as was used for DNA sequencing, using the Phase Genomics HiC Plant Kit version 4.0 (CITE). Samples were sent to the Cornell Biotechnology Resource Center Genomics Facility for 2×150 Paired End sequencing on an Illumina NextSeq 500 instrument. A total of 48.4 Gb of data was generated. The quality control script provided by Phase Genomics was used to assess the HiC data quality.

### Genome size estimate

To generate a histogram of k-mers, Jellyfish version 2.3.0 was used to count all canonical (-C) 21-mers (-m 21) from the PacBio HiFi reads using the command jellyfish count, and a histogram was output using the command jellyfish hist(Marçais & Kingsford 2011). The histogram was fed into the online interface of GenomeScope 2.0 to generate a genome size estimate(Ranallo-Benavidez et al. 2020). A 21-mer was the kmer length recommended for use with the GenomeScope 2.0 program and was not adjusted because we had high coverage and a low error rate.

### Genome assembly

To generate an initial assembly, PacBio CCS Hifi reads were assembled using the *de novo* assembler hifiasm using default parameters(Cheng et al. 2021). Then, in order to improve the genome using proximity information, the HiC data was used to edit the hifiasm assembly using the Juicebox Assembly Tools pipeline(Dudchenko et al.; Durand et al. 2016) with the following steps: (1) the HiC data was aligned to the existing assembly using juicer, (2) the assembly was reordered based on the HiC data using 3D-DNA, and (3) the pseudochromosome boundaries and scaffold orientations were manually edited in Juicebox according to the HiC contact map: regions with inversion errors with “bowtie” motifs were flipped to create a continuous bright band of high contact frequency along the diagonal, and the pseudochromosome boundaries were edited to conform to very clear visually apparent boundaries. BUSCO using the OrthoDB v10 embryophyta dataset was used to assess genome quality (Manni et al. 2021; Kriventseva et al. 2019). A ∼10Mb pseudochromosome containing a large number of putative centromeric repeats and chloroplastic sequences and no BUSCO gene content was assigned a “debris” label (scaffold 1242). The 8 chromosomes were much larger than all other scaffolds and were visually clear in the HiC contact map; these were designated as the chromosomal assembly.

### Genome completeness estimate

In order to assess the completeness of the chromosomal assembly and rule out the possibility of an important quantity of nuclear genome sequence in the remaining scaffolds, we performed a Merqury completeness analysis. First, we ran the included Merqury script best_k.sh to find the best kmer size, which was k=19. Then, in order to generate kmer counts from our PacBio CCS data, we ran meryl v1.3 using the command: meryl count k=19 PacBio.fastq.gz output pacbio.meryl. We then ran the completeness analysis using Merqury v1.3 using the pacbio.meryl file and default parameters.

### Repeat assessment of non-chromosomal sequence

In order to further confirm that the non-chromosomal sequence did not include chromosomal repeats that were excluded from the chromosomal assembly, we ran the Extensive de novo TE Annotator (EDTA) on the data using the parameter “--anno 1” and all default parameters otherwise(Ou et al. 2019).

### Raw Hi-C alignment to non-chromosomal and chromosomal sequence

In order to further confirm that the non-chromosomal sequence did not contain high nuclear DNA content, the raw Hi-C data was aligned to the chromosomal and the non-chromosomal scaffold separately using STAR version 2.7.5a (Dobin et al. 2013). Separate indices were created for the chromosomal scaffolds and the non-chromosomal scaffolds using ‘mode --genomeGenerate’ with default parameters, then STAR was run using parameters ‘--twopassMode Basic --limitOutSJcollapsed 5000000 --limitSjdbInsertNsj 2000000’.

### Genome annotation

A repeat library was made using RepeatModeler with option -LTRStruct(Flynn et al. 2020). Then reads were softmasked using RepeatMasker with option -nolow(SMIT A. F. A 2004) (**Supplementary Table 9**). BRAKER2 version 2.1.6 was run twice, first with protein hints using the OrthoDB v10 plant database as evidence, and then with RNAseq data aligned using STAR version 2.7.5a with ‘--twopassMode Basic’ and default parameters(Brůna et al. 2021; Hoff et al. 2019; Brůna et al. 2020; Buchfink et al. 2015; Iwata & Gotoh 2012). The outputs from the two BRAKER runs were combined using TSEBRA(Gabriel et al. 2021). BUSCO using the OrthoDB v10 embryophyta dataset was used to assess annotation quality (Manni et al. 2021; Kriventseva et al. 2019).

In order to get human-readable gene names, AHRD version 3.3.3 was run on the putative protein sequences using default parameters(Boecker). In order to generate GO terms, InterProScan version 5.55-88.0 was run on the putative protein sequences with the options -f XML --goterms --pathways –iprlookup -t p(Jones et al. 2014). The putative protein sequences were also aligned to the UniRef90 database using Diamond(Buchfink et al. 2021). The XML outputs from InterProScan and Diamond were then fed into BLAST2GO using the options -properties annotation.prop -useobo go.obo -loadblast blastresults.xml - loadips50 ipsout.xml -mapping -annotation -statistics all, which generated GO terms(Götz et al. 2008).

In order to find centromeric repeats, Tandem Repeats Finder was run using the parameters “2 7 7 80 10 50 2000 -h” and the most abundant repeat was assumed to be a partial centromeric repeat. The sequence of the partial centromeric repeat is included in supplementary information (**Supplementary File 2**).

False “annotations” that were annotated by BRAKER from the centromeric repeats near the center of chromosome 4 were manually removed from the dataset by performing a BLAST of the repeat sequence against the amino acid sequences of the annotation, then using seqtk to remove the sequences that came up as BLAST hits. These 221 removed “annotations” did not have BLAST2GO GO terms, and AHRD either marked them as “Unknown protein” or they were missing from the dataset. A full list of the removed sequences is included in supplementary information (**Supplementary File 3**).

### Genome visualization

In order to visualize the distribution of features across the length of the chromosomes, KaryoPloteR was used in R; gff files containing the locations of TEs, centromeric repeats, and gene models were converted to densities using the gffToGRanges() function(Gel & Serra 2017). The HiC contact map was visualized in the Juicebox desktop application version 1.11.08 at the default resolution(Durand et al. 2016).

### Other genome assemblies used for comparative genomics

The Wang et al *Euphorbia lathyris* genome assembly and annotation was accessed through figshare(Mingcheng Wang et al. 2021): https://figshare.com/articles/dataset/High-quality_genome_assembly_of_the_biodiesel_plant_Euphorbia_lathyris/14909913/1 The Liu et al *Hevea brasiliensis* (rubber tree) assembly and annotation was accessed through NCBI(Liu et al. 2020): https://www.ncbi.nlm.nih.gov/assembly/GCA_010458925.1/ The Xu et al wild *Ricinus communis* genome was accessed through oilplantDB(Wei Xu et al. 2021): http://oilplants.iflora.cn/Download/castor_download.html The Bredeson et al *Manihot esculenta* v8.1 genome was accessed through Phytozome: https://phytozome-next.jgi.doe.gov/info/Mesculenta_v8_1

### Macrosynteny analysis

In order to make a multi-genome graphical comparison of synteny, we ran the default version 0.9.3 GENESPACE pipeline, which uses MCScanX and OrthoFinder to get orthogroups within syntenic regions and then projects the position of every orthogroup in the dataset against a single genome(Lovell et al.; Emms & Kelly 2019; Wang et al. 2012).

### Multi-species TE analysis

In order to produce comparable transposable element data across different Euphorbiaceae species’ genomes, we selected only the chromosomes for each species and modeled repeats using an identical pipeline. For each species, a repeat library was made using RepeatModeler with option -LTRStruct(Flynn et al. 2020). Then reads were softmasked using RepeatMasker with default options(SMIT A. F. A 2004) (**Supplementary Tables 10-14**). The TETools script calcDivergenceFromAlign.pl was used to generate the Kimura matrix for each species, then the results of that pipeline were visualized in R using ggplot2(Wickham 2016; Ripley 2001).

### Identifying orthologous genes of interest

In order to produce orthologous groups, we ran OrthoFinder version 2.5.1 using the defaults, including Diamond as the sequence search program, on the protein files from *E. peplus* and the other publicly retrieved Euphorbiaceae genome assemblies(Emms & Kelly 2019; Buchfink et al. 2021). The OrthoFinder species tree was visualized using FigTree v1.4.4. We retrieved the sequences of genes of interest from NCBI and TAIR and ran blastp or tblastn as appropriate with our *E. peplus* proteins file as the query and the parameters -qcov_hsp_perc 80 -evalue 1e-10(Camacho et al. 2009). For our BLAST search for NRPE1, we also ran a BLAST using NRPD1 because the two proteins share a functional domain and we wanted to ensure that we did not misidentify NRPD1 paralogs as a NRPE1 duplication; our NRPE1 paralogs were not a BLAST hit for NRPD1 as a subject. The NRPE1s were aligned using MAFFT v7.453 using the parameters --maxiterate 1000 --localpair and visualized in UniPro UGENE(Katoh & Standley 2013; Okonechnikov et al. 2012). The FastTree NRPE1 phylogeny produced by OrthoFinder was visualized using FigTree v1.4.4.

### Differential gene expression

In order to evaluate differential gene expression between tissues, the results from the STAR alignment were used with the htseq-count script from the HTSeq package to get raw read counts for each RNAseq sample(Anders et al. 2014). Using the DESeq2 package in R, the data with summed counts less than 20 across samples was eliminated, then the DESeq default differential expression analysis was run(Love et al. 2014). A variance stabilizing transformation was applied to the data for PCA visualization. After an initial PCA visualization of all data, one degraded root sample with low read counts was removed and the analysis was re-run **(Supplementary Figure 2, Supplementary Table 3)**. Regularized log-scaled counts of genes of interest were plotted using pheatmap (Kolde).

### eFP Browser

In order to generate an interactive visualization of gene expression across different organs, we first generated transcripts per million (TPM) data from our raw RNAseq reads. GTFTools with argument -l was used to calculate gene length(Li 2018). TPM was then calculated manually in R by dividing the gene length over 1000 to get the length in kb, then dividing the read counts by that number to get reads per kilobase (RPK), then using the prop.table() function to calculate the value of each RPK value as a proportion of the total sum of all RPK values then multiplying by 1,000,000 to get transcripts per million (TPM). A drawing of a *E. peplus* plant including different organs was created in Adobe Illustrator. These data were databased to the Bio-Analytic Resource for Plant Biology (BAR) website as a novel electronic Fluorescent Pictograph (eFP) browser (modified based on code from Winter et al. 2007) where each gene’s TPM can be visualized in each of the six sampled *E. peplus* organs(Winter et al. 2007; Sullivan et al. 2019). The resource is publicly available at https://bar.utoronto.ca/efp_euphorbia/cgi-bin/efpWeb.cgi

### JBrowse

In order to make a publicly accessible visualization of the genome annotation, we implemented a genome web browser in JBrowse version 1.16.11(Buels et al. 2016). The website was certified through Let’s Encrypt(Aas et al. 2019). The resource is publicly available at https://euphorbgenomes.biohpc.cornell.edu/.

### Imaging

Overview plant images were taken with an iPhone 10R. Images of latex dripping were taken with a Canon EOS 80D. Dissecting microscopy images were taken with a Leica M205 FCA stereo microscope with a DMC6200 camera.

## Data availability statement

All sequence data and the completed genome are available through NCBI PRJNA837952 “Euphorbia peplus Genome sequencing and assembly”. All scripts used in the analysis are available on a public Github repository: https://github.com/ariellerjohnson/Euphorbia-peplus-genome-project A plant grown from a seed of the sequenced plant was deposited as a herbarium voucher in the L. H. Bailey Hortorium Herbarium at Cornell University, collection number Ashley Bao AB001. A JBrowse instance of the genome assembly and annotation is publicly available at https://euphorbgenomes.biohpc.cornell.edu/. An eFP Browser instance showing organ-specific gene expression levels is available at https://bar.utoronto.ca/efp_euphorbia/cgi-bin/efpWeb.cgi

## Acknowledgements

We would like to thank Qi Sun of the Cornell University Institute of Biotechnology BioHPC for his generous help with JBrowse and with the BioHPC Docker system. We would like to acknowledge the staff of the Weill Hall growth chambers and the Purple Greenhouse at Cornell for their important assistance with watering and pest maintenance. We would like to thank Peter Fraissinet of the L.H. Bailey Hortorium Herbarium for his generous assistance with the herbarium voucher preparation. We would also like to thank the members of the Frank Lab for helpful comments on this manuscript, as well as Alexandra Bennett of the Moghe Lab for helping with the foundational metabolomic and germplasm sampling work on this project. This work was supported by a Cornell Institute of Biotechnology Seed Grant to G.M. and M.H.F., National Science Foundation Integrative Organismal Systems (NSF IOS) award 1942437 to M.H.F., and Frank lab and Moghe lab startup funds from the College of Agriculture and Life Sciences at Cornell University.

