## Supplemental Figures and Tables for "Chromosome-level genome assembly of *Euphorbia peplus*, a model system for plant latex, reveals that relative lack of Ty3 transposons contributed to its small genome size"

Supplementary Figures and Tables

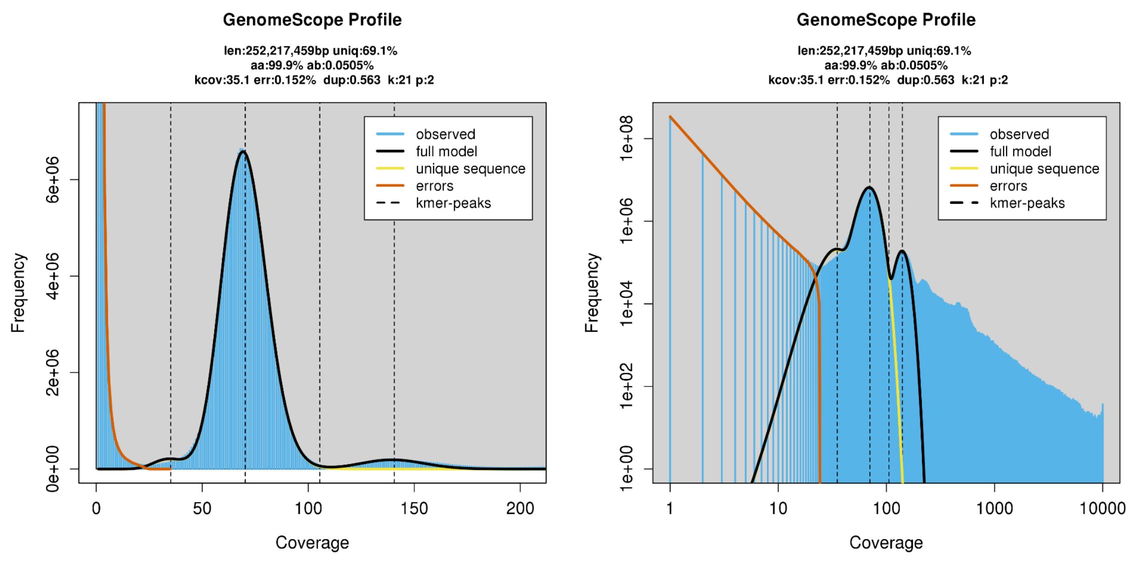

Supplementary Figure 1: GenomeScope2.0 profile of 21-mers in *Euphorbia peplus* raw reads used for genome assembly.

Supplementary Table 1: Genome statistics

| Stat | All scaffolds | 8 chromosomes only |
|---|---|---|
| Total length | 330517859, n = 1242 | 267196017, n = 8 |
| N50 | 31008493, n = 5 | 34601726, n = 4 |
| N60 | 29994419, n = 6 | 31008493, n = 5 |
| N70 | 29603500, n = 7 | 29994419, n = 6 |
| N80 | 29100068, n = 8 | 29603500, n = 7 |
| N90 | 44411, n = 207 | 29100068, n = 8 |
| N100 | 1000, n = 1242 | 29100068, n = 8 |
| Gaps | 239 | 21 |

Supplementary Table 2: BUSCO results for genome assembly

| embryophyta_odb10 (Creation date: 2020-09-10, number of genomes: 50, number of BUSCOs: 1614) | |
|---|---|
| C:98.5%[S:95.5%,D:3.0%],F:0.7%,M:0.8%,n:1614 | |
| 1590 | Complete BUSCOs (C) |
| 1542 | Complete and single-copy BUSCOs (S) |
| 48 | Complete and duplicated BUSCOs (D) |
| 12 | Fragmented BUSCOs (F) |
| 12 | Missing BUSCOs (M) |
| 1614 | Total BUSCO groups searched |

Supplementary Table 3: Merqury completeness results for chromosomal genome only

| assembly | kmer set used for measuring completeness | solid k-mers in the assembly | total solid kmers in the read set | Completeness (%) |
|---|---|---|---|---|
| chromosome_only_genome | all | 182941528 | 184951646 | 98.9132 |

Supplementary Table 4: EDTA results for non-chromosomal scaffolds only

```
Total Sequences: 1234
Total Length: 63212842 bp
Class              Count      bpMasked    %masked
=====              =====      ========    =======
LTR                 --          --          --
   Copia           129        279931      0.44%
   Gypsy            40        134406      0.21%
   unknown          48         85501      0.14%
TIR                 --          --          --
   CACTA             2          6070      0.01%
   Mutator         228         61495      0.10%
nonTIR              --          --          --
   helitron         21         43597      0.07%
                ---------------------------------
   total interspersed 468     611000      0.97%

---------------------------------------------------------
Total              468        611000      0.97%
```

Supplementary Table 5: RNAseq sample metrics

| Sample number | Sample label | Raw read pairs | Number of surviving read pairs after Trimmomatic | Percent of trimmed read pairs uniquely mapped to genome by STAR | Average mapped length |
|---|---|---|---|---|---|
| 1 | fruit_1 | 33435294 | 19121144 | 96.89% | 298.19 |
| 2 | fruit_2 | 22904825 | 12739825 | 97.79% | 298.16 |
| 3 | fruit_3 | 14753156 | 8231045 | 97.74% | 298.01 |
| 4 | inflorescence_1 | 19221750 | 11477518 | 97.25% | 298.18 |
| 5 | inflorescence_2 | 71753516 | 44253871 | 97.15% | 298.17 |
| 6 | inflorescence_3 | 64840066 | 39102768 | 97.30% | 298.02 |
| 7 | root_1 | 14693497 | 8252723 | 97.26% | 298 |
| 8 | (REMOVED root) | 54169 | 31202 | 70.57% | 295.74 |
| 9 | root_3 | 18194801 | 10397279 | 97.19% | 297.99 |
| 10 | young_leaf_1 | 36042628 | 18074168 | 96.29% | 299.30 |
| 11 | young_leaf_2 | 24392458 | 13226057 | 96.02% | 299.26 |
| 12 | young_leaf_3 | 20521163 | 9259620 | 96.62% | 298.81 |
| 13 | stem_1 | 22211664 | 12298940 | 97.31% | 299.32 |
| 14 | stem_2 | 23113957 | 13481043 | 97.41% | 299.10 |
| 15 | stem_3 | 20797223 | 10357684 | 97.10% | 298.91 |
| 16 | mature_leaf_1 | 21174429 | 11968854 | 96.06% | 299.33 |
| 17 | mature_leaf_2 | 19995971 | 10650161 | 97.01% | 299.09 |
| 18 | mature_leaf_3 | 21133127 | 12313830 | 96.26% | 299.51 |

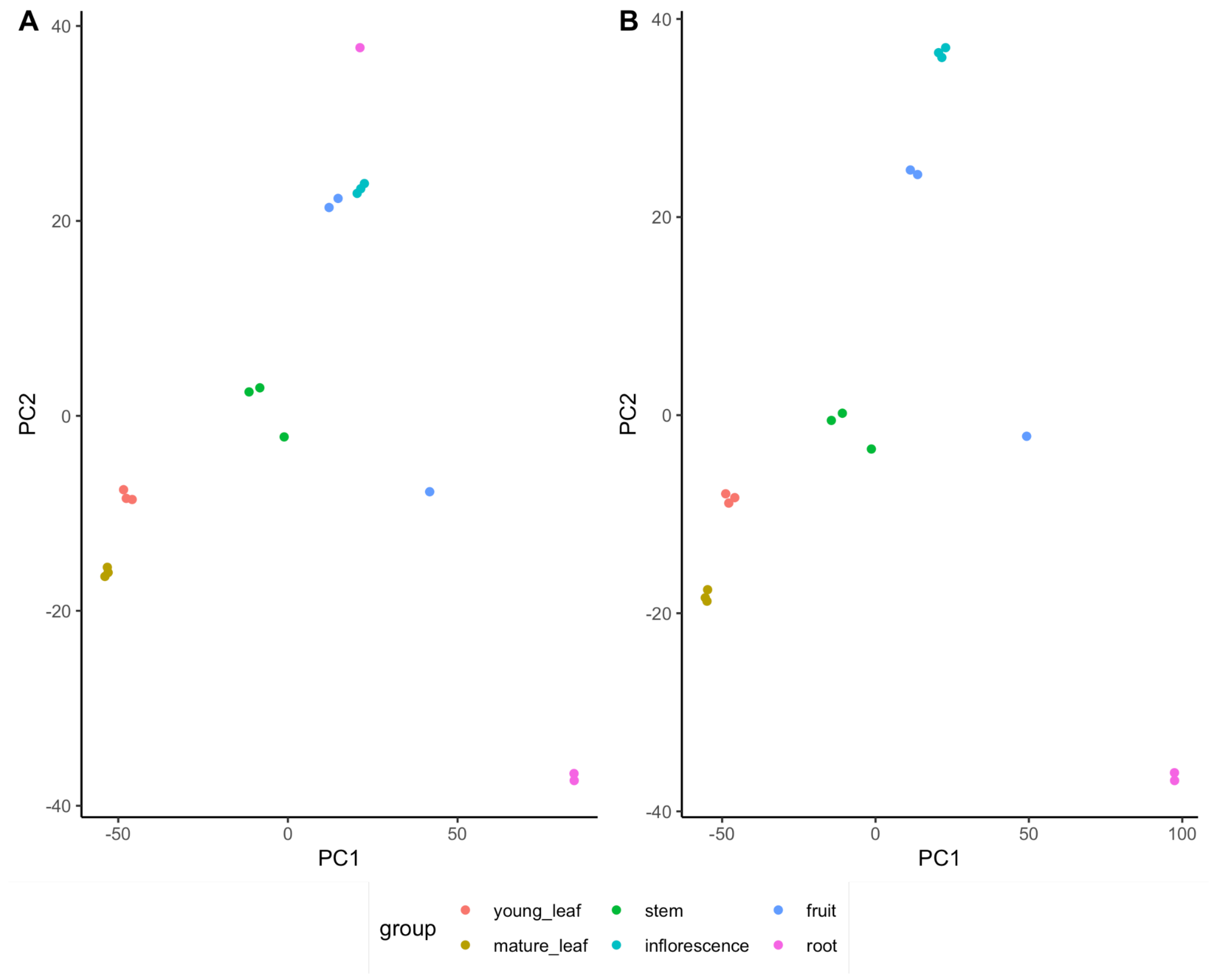

Supplementary Figure 2: PCA plot of variance stabilizing-transformed RNAseq data before and after removal of degraded root sample from RNAseq dataset. A. Before removal of degraded root sample from dataset. B. After removal of degraded root sample from dataset.

Supplementary Table 6: BUSCO results for genome annotation

| | |
|---|---|
| embryophyta_odb10 (Creation date: 2020-09-10, number of genomes: 50, number of BUSCOs: 1614) | |
| C:99.0%[S:90.1%,D:8.9%],F:0.4%,M:0.6%,n:1614 | |
| 1598 | Complete BUSCOs (C) |
| 1454 | Complete and single-copy BUSCOs (S) |
| 144 | Complete and duplicated BUSCOs (D) |
| 6 | Fragmented BUSCOs (F) |

| | |
|---|---|
| 10 | Missing BUSCOs (M) |
| 1614 | Total BUSCO groups searched |

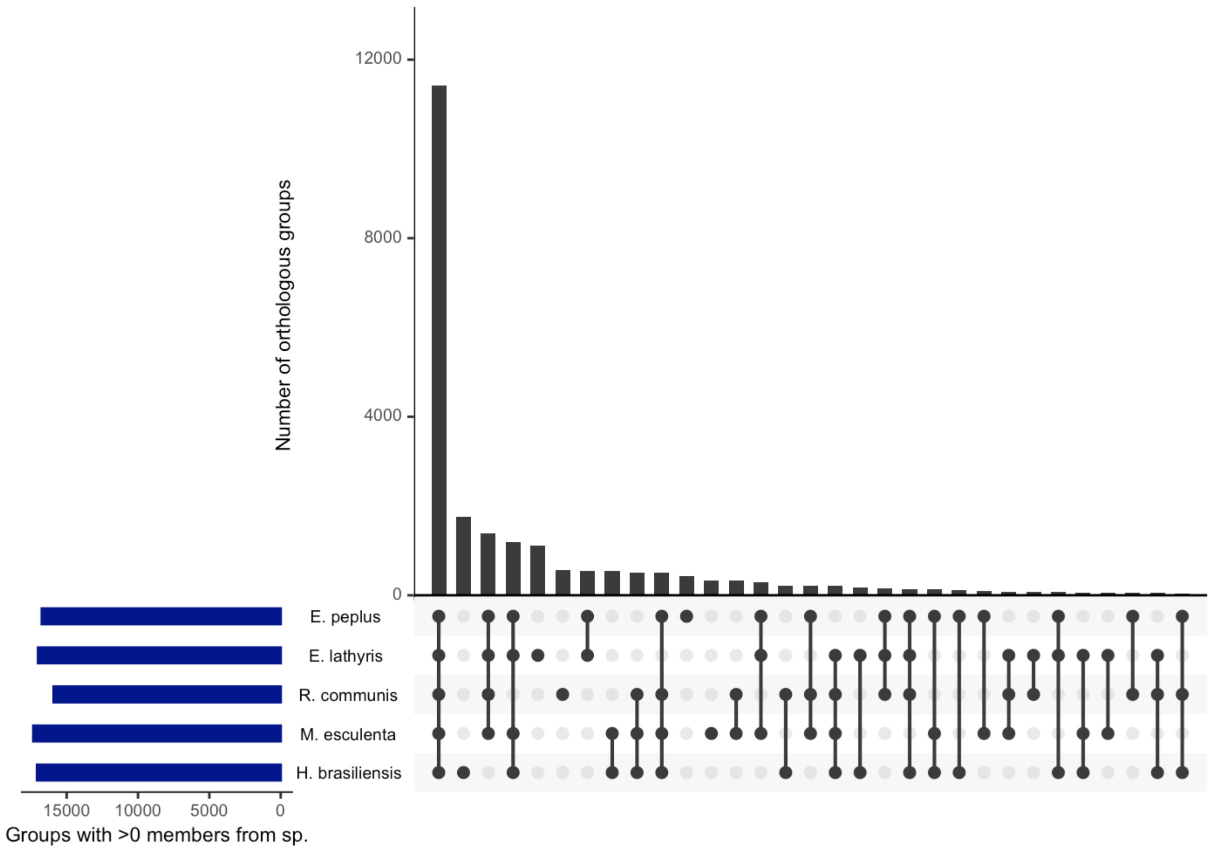

Supplementary Figure 3: UpSet plot of OrthoFinder orthologous groups with members from different combinations of species.

Supplementary Table 7: top BLAST results for *A. thaliana* TE suppression genes of interest

| Gene | qcovs | qseqid | sseqid | pident | length | mismatch | gapopen | evalue |
|---|---|---|---|---|---|---|---|---|
| AGO1 | 89 | Ep_chr6_g16591.t1 | sp\|O04379-2\|AGO1_ARATH | 82.335 | 985 | 159 | 9 | 0.0 |
| AGO9 | 94 | Ep_chr7_g22946.t1 | sp\|Q84VQ0\|AGO9_ARATH | 68.111 | 900 | 264 | 9 | 0.0 |
| ATXR5 | 90 | Ep_chr4_g09628.t1 | sp\|Q8VZJ1- | 61.371 | 321 | 119 | 2 | 9.23e- |

| | | | | | | | | |
|---|---|---|---|---|---|---|---|---|
| | | | 2\|ATXR5_ARATH | | | | | 150 |
| ATXR6 | 95 | Ep_chr4_g09628.t1 | sp\|Q9FNE9\|ATXR6_ARATH | 69.565 | 345 | 94 | 4 | 2.68e-176 |
| CMT2 | 95 | Ep_chr8_g23646.t1 | sp\|Q94F87\|CMT2_ARATH | 50.311 | 805 | 359 | 13 | 0.0 |
| CMT3 | 89 | Ep_chr5_g15899.t1 | sp\|Q94F88\|CMT3_ARATH | 53.611 | 817 | 343 | 11 | 0.0 |
| DDM1 | 100 | Ep_chr6_g20450.t1 | sp\|Q9XFH4\|DDM1_ARATH | 71.466 | 757 | 204 | 8 | 0.0 |
| DRM1 | 93 | Ep_chr6_g16789.t1 | sp\|Q9LXE5\|DRM1L_ARATH | 55.429 | 525 | 219 | 7 | 0.0 |
| DRM2 | 91 | Ep_chr6_g16789.t1 | sp\|Q9M548\|DRM2_ARATH | 56.818 | 528 | 196 | 10 | 0.0 |
| MET1 | 95 | Ep_chr2_g04206.t1 | sp\|P34881\|DNMT1_ARATH | 62.126 | 1505 | 542 | 14 | 0.0 |
| MET2a | 97 | Ep_chr2_g04206.t1 | sp\|O23273-2\|DNMT4_ARATH | 56.577 | 1566 | 607 | 19 | 0.0 |
| NRPD1 | 99 | Ep_chr1_g01405.t1 | sp\|Q9LQ02\|NRPD1_ARATH | 51.131 | 1459 | 661 | 19 | 0.0 |
| NRPE1 | 98 | Ep_chr7_g20683.t1 | sp\|Q5D869\|NRPE1_ARATH | 50.811 | 1972 | 790 | 44 | 0.0 |
| NRPE4 | 99 | Ep_chr4_g10224.t1 | sp\|Q6DBA5\|NRPD4_ARATH | 43.689 | 206 | 103 | 6 | 3.98e-52 |
| NRPE5 | 86 | Ep_chr5_g15247.t1 | sp\|Q9M1J2\|RPE5A_ARATH | 64.286 | 210 | 75 | 0 | 6.92e-105 |
| RDR6 | 100 | Ep_chr4_g12358.t1 | sp\|Q9SG02\|RDR6_ARATH | 67.388 | 1202 | 384 | 6 | 0.0 |

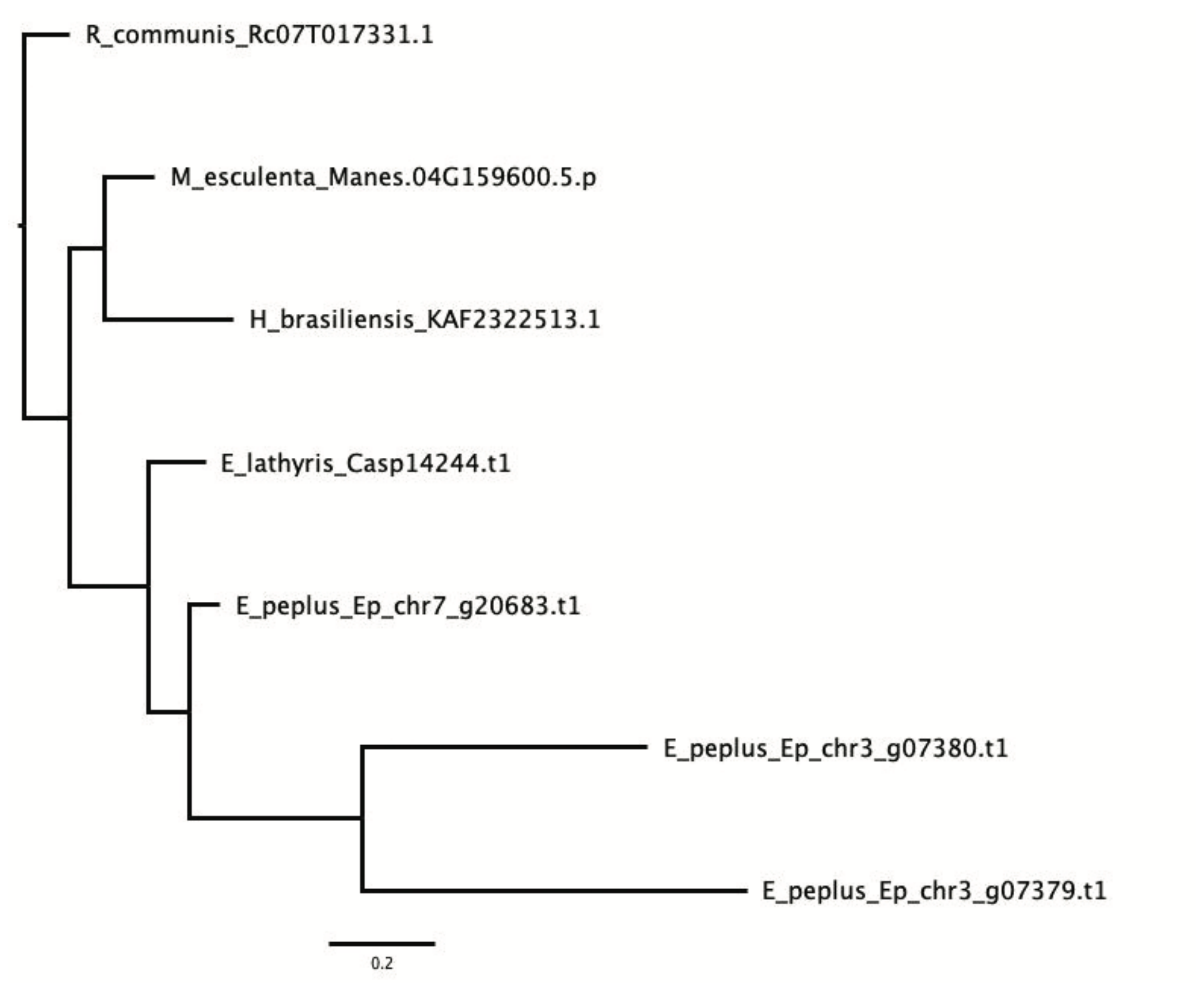

Supplementary Figure 4: Phylogeny of OrthoFinder NRPE1 orthologous group.

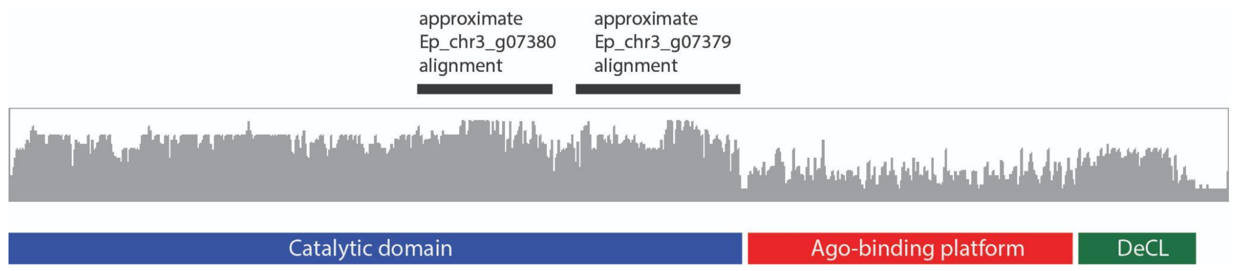

Supplementary Figure 5: Alignment of all genes in the NRPE1 orthologous group, showing NRPE1 regions including non-conserved repetitive Ago-binding platform.

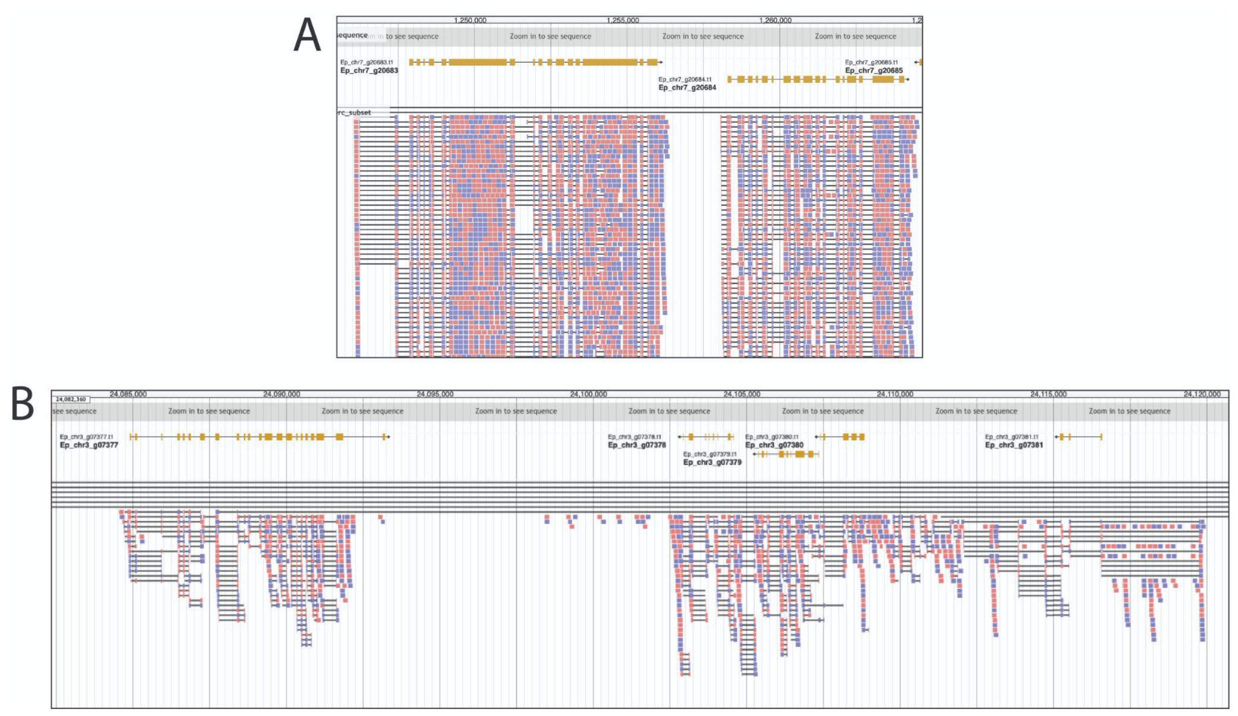

Supplementary Figure 6: Both NRPE paralogs in *E. peplus* show some expression, but the Chromosome 7 ortholog is more highly expressed. A. JBrowse screenshot of Ep_chr7_20683 and a neighboring gene. B. JBrowse screenshot of Ep_chr3_07379 and Ep_chr3_07380 and the surrounding area.

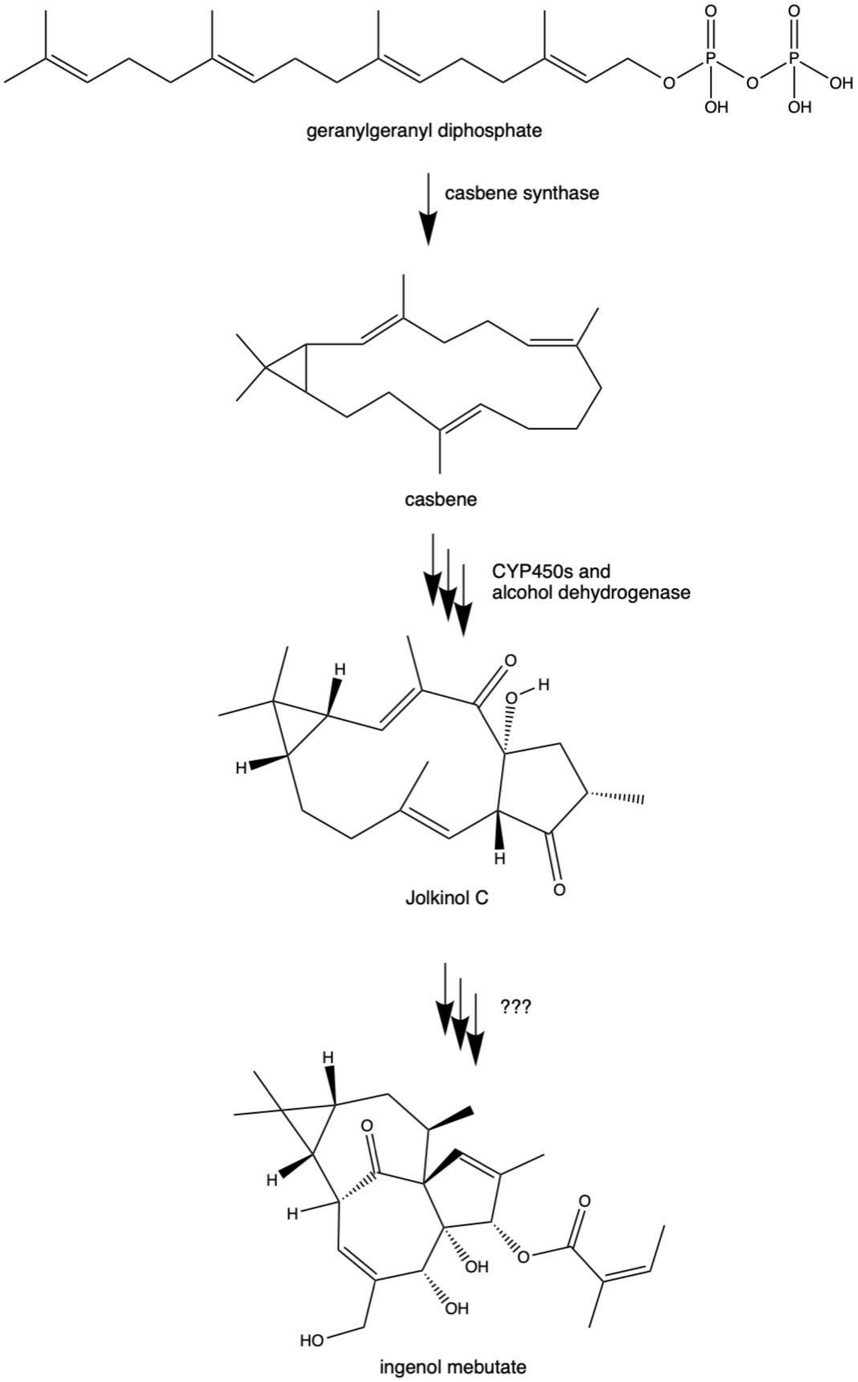

Supplementary Figure 7: simplified ingenol mebutate biosynthetic pathway (modified from Czechowski et al 2022 and Bergman et al 2019).

Supplementary Table 8: Genes in putative diterpene biosynthetic gene cluster (genes below bold line are displayed in Figure 5). CYP names are following Czechowski et al 2022, genes unlikely/unknown to be involved in diterpene synthesis are in parenthesis. CYPs listed by name (e.g. "CYP71D627") without other text were confirmed by BLAST to be the same genes as the sequences deposited by Czechowski et al 2022.

| Gene ID | Gene name | BAC ID from Czechowski et al 2022 | Functionally characterized for *Euphorbia*? How? |
|---|---|---|---|
| Ep_chr6_g18767 | (Proteasome subunit alpha type-2) | B24L04 | |
| Ep_chr6_g18768 | (ATP-dependent rRNA helicase RRP3) | B24L04 | |
| Ep_chr6_g18769 | (unknown protein) | | |
| Ep_chr6_g18770 | (unknown protein) | | |
| Ep_chr6_g18771 | (Mitochondrial transcription termination factor family protein) | | |
| Ep_chr6_g18772 | (mercaptopyruvate sulfurtransferase 1) | B24L04 | |
| Ep_chr6_g18773 | (Sec14p-like phosphatidylinositol transfer family protein) | | |
| Ep_chr6_g18774 | (unknown protein) | | |
| Ep_chr6_g18775 | (S-adenosyl-L-methionine-dependent methyltransferases superfamily protein) | | |
| Ep_chr6_g18776 | (Zinc-binding dehydrogenase family protein) | | |
| Ep_chr6_g18777 | NAD(P)-linked oxidoreductase superfamily protein | | |
| Ep_chr6_g18778 | 2-oxoglutarate (2OG) and Fe(II)-dependent oxygenase superfamily protein | | |
| Ep_chr6_g18779 | Other CYP450–CYP726A9 | B24L04 | |
| Ep_chr6_g18780 | CYP71D627 | B24L04, H22G04 | |
| Ep_chr6_g18781 | 2-oxoglutarate (2OG) and Fe(II)-dependent oxygenase superfamily protein | H22G04 | |

| | | |
|---|---|---|
| Ep_chr6_g18782 | Gibberellin 20 oxidase 2 | |
| Ep_chr6_g18783 | 2-oxoglutarate (2OG) and Fe(II)-dependent oxygenase superfamily protein | |
| Ep_chr6_g18784 | 2-oxoglutarate-dependent dioxygenase DAO | H22G04 |
| Ep_chr6_g18785 | (Protein kinase superfamily protein with octicosapeptide/Phox/Bem1p domain) | H22G04 |
| Ep_chr6_g18786 | (Protein kinase superfamily protein with octicosapeptide/Phox/Bem1p domain) | H22G04 |
| Ep_chr6_g18787 | (GRAS family transcription factor) | |
| Ep_chr6_g18788 | (nuclear factor Y, subunit C2) | |
| Ep_chr6_g18789 | (SKP1-like 12) | |
| Ep_chr6_g18790 | 2,4-dienoyl-CoA reductase_1 | |
| Ep_chr6_g18791 | 2,4-dienoyl-CoA reductase_2 | |
| Ep_chr6_g18792 | 2,4-dienoyl-CoA reductase_3 | |
| Ep_chr6_g18793 | (pseudouridine synthase family protein) | |
| Ep_chr6_g18794 | other CYP450_1–CYP76F147 | B17P14, H22G04 |
| Ep_chr6_g18795 | (ATP-dependent zinc metalloprotease) | |
| Ep_chr6_g18796 | (mis-annotated part of following gene) | B17P14, H22G04 |
| Ep_chr6_g18797 | (unknown protein_1) | B17P14, H22G04 |
| Ep_chr6_g18798 | (phosphoglycerate kinase family protein) | B17P14, H22G04 |
| Ep_chr6_g18799 | (phosphoglycerate kinase) | B17P14, H22G04 |
| Ep_chr6_g18800 | CYP726A46 | B17P14 |
| Ep_chr6_g18801 | myrcene synthase_1 | B17P14 |
| Ep_chr6_g18802 | CYP726A38 | B17P14 |

| | | | |
|---|---|---|---|
| Ep_chr6_g18803 | myrcene synthase_2 | B17P14 | |
| Ep_chr6_g18804 | myrcene synthase_3 | B17P14 | |
| Ep_chr6_g18805 | myrcene synthase_4 | B17P14 | |
| Ep_chr6_g18806 | other CYP450_2–CYP80C15 | B17P14 | |
| Ep_chr6_g18807 | myrcene synthase_5 | B17P14 | |
| Ep_chr6_g18808 | CYP726A39 | B17P14 | |
| Ep_chr6_g18809 | CYP726A45 | B17P14 | |
| Ep_chr6_g18810 | other CYP450_3–CYP726A5 | B17P14 | |
| Ep_chr6_g18811 | CYP71D369 | B17P14 | |
| Ep_chr6_g18812 | other CYP450_4–CYP726A6 | B17P14 | |
| Ep_chr6_g18813 | (NAD(P)-binding Rossmann-fold superfamily protein) | B17P14 | |
| Ep_chr6_g18814 | CYP726A42 | B17P14 | |
| Ep_chr6_g18815 | CYP726A19 | B17P14 | |
| Ep_chr6_g18816 | casbene synthase | B17P14 | VIGS in *Euphorbia peplus* (Czechowski et al 2022) |
| Ep_chr6_g18817 | dienoyl-CoA isomerase_1–"crotonase 1" in Czechowski et al? | B17P14? | |
| Ep_chr6_g18818 | dienoyl-CoA isomerase_2 –"crotonase 1" in Czechowski et al? | B17P14? | |
| Ep_chr6_g18819 | dienoyl-CoA isomerase_3 –"crotonase 1" in Czechowski et al? | B17P14? | |
| Ep_chr6_g18820 | (unknown protein_2) | | |
| Ep_chr6_g18821 | (polyketide cyclase/dehydrase and lipid transport superfamily protein) | | |
| Ep_chr6_g18822 | CYP726A43 | H07C08 | *N. benthamiana* and *S. cerevisiae* heterologous expression of *Euphorbia lathyris* CYP726A27 (Luo |

| | | | |
|---|---|---|---|
| | | | et al 2016) |
| Ep_chr6_g18823 | CYP726A3 | H07C08 | |
| Ep_chr6_g18824 | CYP726A4 | P10A24 | |
| Ep_chr6_g18825 | carboxylesterase | P10A24 | |
| Ep_chr6_g18826 | CYP71D365 | P10A24 | VIGS in *Euphorbia peplus* (Czechowski et al 2022); *N. benthamiana* and *S. cerevisiae* heterologous expression of *Euphorbia lathyris* CYP71D445 (Luo et al 2016) |
| Ep_chr6_g18827 | CYP726A40 | P10A24 | |
| Ep_chr6_g18828 | CYP726A44 | P10A24 | |
| Ep_chr6_g18829 | dienoyl-CoA isomerase_4 – crotonase 2 | P10A24 | |
| Ep_chr6_g18830 | CYP71D367 | P10A24 | |
| Ep_chr6_g18831 | CYP726A41 | P10A24 | |
| Ep_chr6_g18832 | (unknown protein_3) | P10A24 | |
| Ep_chr6_g18833 | (Gag-Pol polyprotein) | P10A24 | |
| Ep_chr6_g18834 | CYP71D625 | P10A24 | |
| Ep_chr6_g18835 | other CYP450_5 | | |
| Ep_chr6_g18836 | other CYP450_6 | | |
| Ep_chr6_g18837 | other CYP450_7 | | |
| Ep_chr6_g18838 | CYP71D626 | P10A24 | |
| Ep_chr6_g18839 | CYP71D360 | P10A24 | |

Supplementary Table 9: *E. peplus* all scaffolds RepeatMasker results

```
==================================================
file name: Euphorbia_peplus.fa
sequences:        1242
total length:  330517859 bp  (330398359 bp excl N/X-runs)
GC level:        36.11 %
bases masked:  190579965 bp ( 57.66 %)
==================================================
               number of      length   percentage
               elements*    occupied  of sequence
--------------------------------------------------
Retroelements        53036     64311228 bp   19.46 %
   SINEs:              286      2574730 bp    0.78 %
   Penelope              0            0 bp    0.00 %
   LINEs:            11815      6881222 bp    2.08 %
    CRE/SLACS            0            0 bp    0.00 %
     L2/CR1/Rex        285        67176 bp    0.02 %
     R1/LOA/Jockey     341      2718854 bp    0.82 %
     R2/R4/NeSL          0            0 bp    0.00 %
     RTE/Bov-B        1829       276776 bp    0.08 %
     L1/CIN4          9229      3794464 bp    1.15 %
   LTR elements:     40935     54855276 bp   16.60 %
     BEL/Pao             0            0 bp    0.00 %
     Ty1/Copia       17531     36556366 bp   11.06 %
     Gypsy/DIRS1     12134     14427075 bp    4.36 %
       Retroviral        0            0 bp    0.00 %

DNA transposons       8445      4135604 bp    1.25 %
   hobo-Activator     1490       546064 bp    0.17 %
   Tc1-IS630-Pogo      895       367698 bp    0.11 %
   En-Spm                0            0 bp    0.00 %
   MuDR-IS905            0            0 bp    0.00 %
   PiggyBac              0            0 bp    0.00 %
   Tourist/Harbinger  1917       702225 bp    0.21 %
   Other (Mirage,        0            0 bp    0.00 %
    P-element, Transib)

Rolling-circles       4366      2374903 bp    0.72 %

Unclassified:       241792     86528019 bp   26.18 %

Total interspersed repeats:   154974851 bp   46.89 %

Small RNA:            8959     27599976 bp    8.35 %

Satellites:           1103      5612899 bp    1.70 %
Simple repeats:        398        17336 bp    0.01 %
Low complexity:          0            0 bp    0.00 %
```

```
==================================================

* most repeats fragmented by insertions or deletions
  have been counted as one element

RepeatMasker version 4.1.2-p1 , default mode

run with rmblastn version 2.11.0+
The query was compared to classified sequences in "Euphorbia_peplus-families.fa"
FamDB:
```

Supplementary Table 10: *E. peplus* chromosomes RepeatMasker results

```
==================================================
file name: E_peplus.fa
sequences:            8
total length:  267196017 bp  (267185517 bp excl N/X-runs)
GC level:         35.22 %
bases masked:  129714136 bp ( 48.55 %)
==================================================
               number of      length   percentage
               elements*    occupied  of sequence
--------------------------------------------------
Retroelements        47754     59183173 bp   22.15 %
   SINEs:                0            0 bp    0.00 %
   Penelope              0            0 bp    0.00 %
   LINEs:            10633      3143544 bp    1.18 %
    CRE/SLACS            0            0 bp    0.00 %
     L2/CR1/Rex        165        35988 bp    0.01 %
     R1/LOA/Jockey       0            0 bp    0.00 %
     R2/R4/NeSL          0            0 bp    0.00 %
     RTE/Bov-B        2349       320544 bp    0.12 %
     L1/CIN4          8119      2787012 bp    1.04 %
   LTR elements:     37121     56039629 bp   20.97 %
     BEL/Pao             0            0 bp    0.00 %
     Ty1/Copia       17494     39345359 bp   14.73 %
     Gypsy/DIRS1      8739     12729617 bp    4.76 %
       Retroviral        0            0 bp    0.00 %

DNA transposons       8763      3805128 bp    1.42 %
   hobo-Activator      963       474623 bp    0.18 %
   Tc1-IS630-Pogo      866       370951 bp    0.14 %
   En-Spm                0            0 bp    0.00 %
   MuDR-IS905            0            0 bp    0.00 %
   PiggyBac              0            0 bp    0.00 %
```

```
  Tourist/Harbinger  3112       815771 bp    0.31 %
  Other (Mirage,        0            0 bp    0.00 %
   P-element, Transib)

Rolling-circles       4377      2074134 bp    0.78 %

Unclassified:       233798     61053976 bp   22.85 %

Total interspersed repeats:   124042277 bp   46.42 %

Small RNA:             721       363226 bp    0.14 %

Satellites:              0            0 bp    0.00 %
Simple repeats:      60110      2625352 bp    0.98 %
Low complexity:      12542       609147 bp    0.23 %
==================================================

* most repeats fragmented by insertions or deletions
  have been counted as one element

RepeatMasker version 4.1.2-p1 , default mode

run with rmblastn version 2.11.0+
The query was compared to classified sequences in "E_peplus-families.fa"
FamDB:
```

Supplementary Table 11: *E. lathyris* chromosomes RepeatMasker results

```
==================================================
file name: E_lathyris.fa
sequences:            10
total length:  986794769 bp  (986783869 bp excl N/X-runs)
GC level:         36.83 %
bases masked:  766851039 bp ( 77.71 %)
==================================================
               number of      length   percentage
               elements*    occupied  of sequence
--------------------------------------------------
Retroelements       474794    548252351 bp   55.56 %
   SINEs:                0            0 bp    0.00 %
   Penelope              0            0 bp    0.00 %
   LINEs:            66912     54340451 bp    5.51 %
    CRE/SLACS            0            0 bp    0.00 %
     L2/CR1/Rex          0            0 bp    0.00 %
     R1/LOA/Jockey     270        51422 bp    0.01 %
     R2/R4/NeSL          0            0 bp    0.00 %
```

```
    RTE/Bov-B         228        47023 bp    0.00 %
    L1/CIN4         65851     54074699 bp    5.48 %
   LTR elements:    407882    493911900 bp   50.05 %
    BEL/Pao            0            0 bp    0.00 %
    Ty1/Copia      204376    253103775 bp   25.65 %
    Gypsy/DIRS1    127814    205514331 bp   20.83 %
      Retroviral       0            0 bp    0.00 %

DNA transposons      33207     19296054 bp    1.96 %
   hobo-Activator     6296      2394483 bp    0.24 %
   Tc1-IS630-Pogo     3352       721126 bp    0.07 %
   En-Spm              0            0 bp    0.00 %
   MuDR-IS905          0            0 bp    0.00 %
   PiggyBac            0            0 bp    0.00 %
   Tourist/Harbinger  2109       966565 bp    0.10 %
   Other (Mirage,       0            0 bp    0.00 %
    P-element, Transib)

Rolling-circles       9201      3441220 bp    0.35 %

Unclassified:       730609    187474123 bp   19.00 %

Total interspersed repeats:   755022528 bp   76.51 %

Small RNA:           1335       296840 bp    0.03 %

Satellites:            33        65451 bp    0.01 %
Simple repeats:     142306      6661152 bp    0.68 %
Low complexity:      26256      1363848 bp    0.14 %
==================================================

* most repeats fragmented by insertions or deletions
  have been counted as one element

RepeatMasker version 4.1.2-p1 , default mode

run with rmblastn version 2.11.0+
The query was compared to classified sequences in "E_lathyris-families.fa"
FamDB:
```

Supplementary Table 12: *H. brasiliensis* chromosomes RepeatMasker results

```
==================================================
file name: H_brasiliensis.fa
sequences:            18
total length: 1441955164 bp  (1440424564 bp excl N/X-runs)
```

GC level: 33.84 %
bases masked: 1087871690 bp ( 75.44 %)

| | number of elements* | length occupied | percentage of sequence |
|---|---|---|---|
| Retroelements | 617134 | 789499983 bp | 54.75 % |
| SINEs: | 0 | 0 bp | 0.00 % |
| Penelope | 0 | 0 bp | 0.00 % |
| LINEs: | 28033 | 16033028 bp | 1.11 % |
| CRE/SLACS | 0 | 0 bp | 0.00 % |
| L2/CR1/Rex | 0 | 0 bp | 0.00 % |
| R1/LOA/Jockey | 0 | 0 bp | 0.00 % |
| R2/R4/NeSL | 0 | 0 bp | 0.00 % |
| RTE/Bov-B | 13936 | 10408242 bp | 0.72 % |
| L1/CIN4 | 13475 | 5492488 bp | 0.38 % |
| LTR elements: | 589101 | 773466955 bp | 53.64 % |
| BEL/Pao | 0 | 0 bp | 0.00 % |
| Ty1/Copia | 215517 | 250019618 bp | 17.34 % |
| Gypsy/DIRS1 | 297464 | 484733236 bp | 33.62 % |
| Retroviral | 0 | 0 bp | 0.00 % |
| DNA transposons | 12875 | 9436633 bp | 0.65 % |
| hobo-Activator | 5636 | 3758509 bp | 0.26 % |
| Tc1-IS630-Pogo | 0 | 0 bp | 0.00 % |
| En-Spm | 0 | 0 bp | 0.00 % |
| MuDR-IS905 | 0 | 0 bp | 0.00 % |
| PiggyBac | 0 | 0 bp | 0.00 % |
| Tourist/Harbinger | 0 | 0 bp | 0.00 % |
| Other (Mirage, P-element, Transib) | 0 | 0 bp | 0.00 % |
| Rolling-circles | 3656 | 2163645 bp | 0.15 % |
| Unclassified: | 614549 | 269790264 bp | 18.71 % |
| Total interspersed repeats: | | 1068726880 bp | 74.12 % |
| Small RNA: | 0 | 0 bp | 0.00 % |
| Satellites: | 714 | 135122 bp | 0.01 % |
| Simple repeats: | 318295 | 13778849 bp | 0.96 % |
| Low complexity: | 57975 | 3067194 bp | 0.21 % |

* most repeats fragmented by insertions or deletions have been counted as one element

RepeatMasker version 4.1.2-p1 , default mode

run with rmblastn version 2.11.0+
The query was compared to classified sequences in "H_brasiliensis-families.fa"
FamDB:

Supplementary Table 13: *M. esculenta* chromosomes RepeatMasker results

```
==================================================
file name: M_esculenta.fa
sequences:            18
total length:  633922132 bp  (631606260 bp excl N/X-runs)
GC level:         37.43 %
bases masked:  433480988 bp ( 68.38 %)
==================================================
               number of      length   percentage
               elements*    occupied  of sequence
--------------------------------------------------
Retroelements       255297    310386046 bp   48.96 %
   SINEs:                0            0 bp    0.00 %
   Penelope             43        27008 bp    0.00 %
   LINEs:            13810      6420213 bp    1.01 %
    CRE/SLACS            0            0 bp    0.00 %
     L2/CR1/Rex          0            0 bp    0.00 %
     R1/LOA/Jockey       0            0 bp    0.00 %
     R2/R4/NeSL          0            0 bp    0.00 %
     RTE/Bov-B        1132       244541 bp    0.04 %
     L1/CIN4         12635      6148664 bp    0.97 %
   LTR elements:    241487    303965833 bp   47.95 %
     BEL/Pao             0            0 bp    0.00 %
     Ty1/Copia       53182     41451773 bp    6.54 %
     Gypsy/DIRS1    142159    222263321 bp   35.06 %
       Retroviral      810       334790 bp    0.05 %

DNA transposons       8800      5776405 bp    0.91 %
   hobo-Activator     3005      1532524 bp    0.24 %
   Tc1-IS630-Pogo        0            0 bp    0.00 %
   En-Spm                0            0 bp    0.00 %
   MuDR-IS905            0            0 bp    0.00 %
   PiggyBac              0            0 bp    0.00 %
   Tourist/Harbinger   450       290030 bp    0.05 %
   Other (Mirage,        0            0 bp    0.00 %
    P-element, Transib)

Rolling-circles       1174       720821 bp    0.11 %

Unclassified:       296032    108671704 bp   17.14 %
```

```
Total interspersed repeats:   424834155 bp   67.02 %

Small RNA:            533       268129 bp    0.04 %

Satellites:             0            0 bp    0.00 %
Simple repeats:     136526      5930092 bp    0.94 %
Low complexity:      32594      1727791 bp    0.27 %
==================================================

* most repeats fragmented by insertions or deletions
  have been counted as one element

RepeatMasker version 4.1.2-p1 , default mode

run with rmblastn version 2.11.0+
The query was compared to classified sequences in "M_esculenta-families.fa"
FamDB:
```

Supplementary Table 14: *R. communis* chromosomes RepeatMasker results

```
==================================================
file name: R_communis.fa
sequences:            10
total length:  328026558 bp  (328017246 bp excl N/X-runs)
GC level:         33.10 %
bases masked:  190167804 bp ( 57.97 %)
==================================================
               number of      length   percentage
               elements*    occupied  of sequence
--------------------------------------------------
Retroelements       111360    116207587 bp   35.43 %
   SINEs:              210        45117 bp    0.01 %
   Penelope              0            0 bp    0.00 %
   LINEs:             1956       822827 bp    0.25 %
    CRE/SLACS            0            0 bp    0.00 %
     L2/CR1/Rex          0            0 bp    0.00 %
     R1/LOA/Jockey     470       198767 bp    0.06 %
     R2/R4/NeSL          0            0 bp    0.00 %
     RTE/Bov-B         276        49821 bp    0.02 %
     L1/CIN4          1210       574239 bp    0.18 %
   LTR elements:    109194    115339643 bp   35.16 %
     BEL/Pao             0            0 bp    0.00 %
     Ty1/Copia       19761     23238317 bp    7.08 %
     Gypsy/DIRS1     61659     65710364 bp   20.03 %
       Retroviral        0            0 bp    0.00 %
```

```
DNA transposons       8882      5118037 bp    1.56 %
   hobo-Activator     1053       701122 bp    0.21 %
   Tc1-IS630-Pogo      655       236476 bp    0.07 %
   En-Spm                0            0 bp    0.00 %
   MuDR-IS905            0            0 bp    0.00 %
   PiggyBac              0            0 bp    0.00 %
   Tourist/Harbinger    59        41033 bp    0.01 %
   Other (Mirage,        0            0 bp    0.00 %
    P-element, Transib)

Rolling-circles        454       337319 bp    0.10 %

Unclassified:       176520     63167632 bp   19.26 %

Total interspersed repeats:   184493256 bp   56.24 %

Small RNA:            1009       409402 bp    0.12 %

Satellites:            303       101836 bp    0.03 %
Simple repeats:      92559      3800548 bp    1.16 %
Low complexity:      21030      1025443 bp    0.31 %
==================================================

* most repeats fragmented by insertions or deletions
  have been counted as one element

RepeatMasker version 4.1.2-p1 , default mode

run with rmblastn version 2.11.0+
The query was compared to classified sequences in "R_communis-families.fa"
FamDB:
```
